# Metabolic reinvigoration of NK cells by IL-21 enhances immunotherapy against MHC-I deficient solid tumors

**DOI:** 10.1101/2025.03.19.643961

**Authors:** Yi Wang, Chao Huang, Guoxin Cai, Massimo Andreatta, Armand Kurum, Yang Zhao, Bing Feng, Min Gao, Santiago J. Carmona, Zhan Zhou, Cheng Sun, Yugang Guo, Li Tang

**Affiliations:** Institute of Bioengineering, École Polytechnique Fédérale de Lausanne (EPFL), 1015 Lausanne, Switzerland; Institute of Materials Science & Engineering, EPFL, 1015 Lausanne, Switzerland; Institute of Immunology, School of Life Sciences, University of Science and Technology of China, Hefei, 230027, China; National Key Laboratory of Advanced Drug Delivery and Release Systems, Zhejiang University, 310058 Hangzhou, China; Institute of Drug Metabolism and Pharmaceutical Analysis, College of Pharmaceutical Sciences, Zhejiang University, 310058 Hangzhou, China; Department of Oncology, University of Lausanne, 1011 Lausanne, Switzerland; Swiss Institute of Bioinformatics, Lausanne, Switzerland

**Keywords:** Interleukin-21, NK cell therapy, MHC-I deficient tumor, NK cell exhaustion, glycolysis, LDHA, cancer immunotherapy, Immunometabolism

## Abstract

Natural killer (NK) cells, a type of potent cytotoxic lymphocytes, are particularly promising for the treatment of cancers that lose or downregulate major histocompatibility complex class I (MHC-I) expression to evade T cell-mediated immunotherapy. However, the hostile and immune suppressive tumor microenvironment (TME) greatly hinders the function of tumor-infiltrating NK cells limiting the therapeutic efficacy against solid tumors. Here, we show that a fusion protein of interleukin-21 (IL-21−Fc), as a direct *in vivo* intervention, can safely and effectively reprogram NK cell metabolism and enhance their effector function in the TME. Our research demonstrates that combining IL-21−Fc with IL-15 superagonist (IL-15SA) or NK cell transfer leads to the eradication of MHC-I-deficient tumors and confers durable protection in syngeneic and xenograft tumor models. Mechanistically, we uncover that IL-21−Fc enhances NK cell effector function by upregulating glycolysis in a lactate dehydrogenase A (LDHA)-dependent manner. These findings not only underscore the considerable potential of IL-21−Fc as an in vivo therapeutic intervention to bolster NK cell-based immunotherapy, but also unveil an innovative strategy of metabolic reprogramming for NK cell rejuvenation within tumors.

**GRAPHICAL ABSTRACT:** 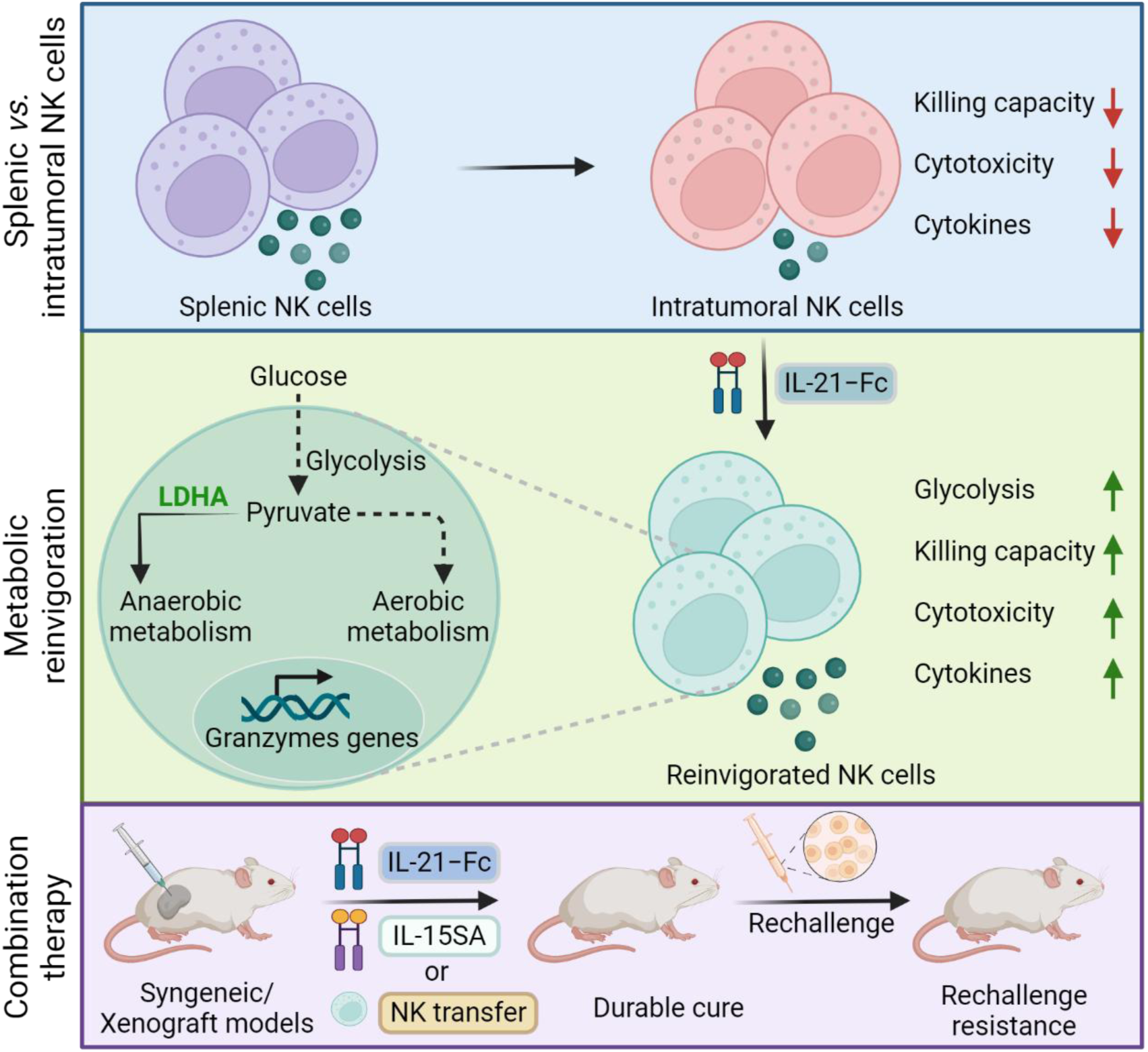

**HIGHLIGHTS:**

- NK cells display functional exhaustion in MHC-I deficient solid tumors
- IL-21−Fc as an in vivo-applicable and safe immunotherapy reinvigorates intratumoral NK cells for enhanced effector function
- IL-21−Fc enhances NK cell function by elevating glycolysis in a LDHA-dependent manner
- Combining IL-21−Fc with low-dose IL-15SA or NK cell transfer eradicates syngeneic and xenografted solid tumors

## INTRODUCTION

Despite the great success of current cancer immunotherapies, the majority of patients fail to respond to immune-related treatments, such as immune checkpoint blockade (ICB)^1,2^. Various mechanisms allow cancer to evade immune surveillance and hinder immunotherapies. Notably, cancer cells frequently downregulate MHC-I expression on their surface to evade recognition and destruction by cytotoxic T cells, a tactic often observed in patients with advanced tumors^3-6^. In addition to T cell-centric immunotherapy, NK cells emerge as potent cytotoxic immune cells capable of recognizing and eliminating cancer cells independent of MHC-I through the dynamic expression of activating and inhibitory receptors^7-10^. Importantly, tumor cells lacking MHC-I become more vulnerable to NK cell-mediated cytotoxicity, rendering NK cell-based immunotherapy a promising alternative to T-cell approaches^11-13^.

Clinical studies utilizing NK cell-based immunotherapy have demonstrated promising results in treating patients with hematological malignancies, showcasing enhanced efficacy and safety^14-17^. However, the hostile TME of solid tumors greatly impedes the antitumor activity of NK cell-based immunotherapy^9,18^. The tumor-infiltrating NK cells often exhibit exhaustion with reduced function in advanced cancers^19-21^. Various intrinsic and extrinsic factors within the TME contribute to impairing their effector function^22,23^. Accumulating evidence suggests that cellular metabolism, including glycolysis and oxidative phosphorylation (OXPHOS), plays a pivotal role in regulating immune cell fate and function^24-27^. Nutrient deprivation in the TME restricts the metabolic activity of NK cells, thereby compromising their effector function^28,29^. Therefore, reprogramming the metabolism of NK cells presents a promising strategy to rejuvenate exhausted NK cells in the TME.

Cytokines are important adjuvant therapeutic agents to promote activation, function, and survival of NK cells for cancer immunotherapy^28,30-33^. Interleukin-21 (IL-21), a cytokine that belongs to the common cytokine receptor gamma-chain family (γ_c_), has been used extensively for *ex vivo* activation of NK cell^34,35^. Notably, strategies such as feeder cells expressing membrane-bound IL-21 have been employed to expand NK cells prior to adoptive transfer therapy, enhancing their cytotoxic capabilities by inducing a shift toward a Warburg metabolism^36^. Furthermore, IL-21 was found to reverse PD-1^+^Tim-3^+^ exhausted NK cell function, particularly in combination with monocyte-based vaccination therapy^19^. However, the feasibility and mechanism underlying the *in vivo* application of IL-21 as a supportive agent for NK cell-based immunotherapy remain unclear.

In the present study, we show that recombinant fusion proteins of IL-21 and Fc (mouse and human) effectively restore metabolic activity and effector function of exhausted mouse and human NK cells. When combined with IL-15SA, a well-established cytokine intervention for augmenting NK cell expansion, or adoptive transfer of NK cells^37,38^, IL-21−Fc eradicated MHC-I-deficient solid tumors in a fraction of treated mice and induced durable protection against rechallenge of the corresponding tumor cells. Furthermore, we elucidate that IL-21−Fc enhances NK cell effector function by elevating glycolysis in a LDHA-dependent manner. These results underscore the potential of metabolic reprogramming of NK cells through the in *vivo* application of cytokines as a promising therapeutic strategy to bolster NK cell-based immunotherapy for cancer.

## RESULTS

### Tumor-infiltrating NK cells are functionally impaired

To assess the function of NK cells within the TME, we established a subcutaneous (s.c.) MHC-I-deficient tumor model by inoculating BALB/c mice with CT26_β2m^-/-^ murine colorectal carcinoma cells (8 × 10^5^), which did not respond to cytotoxic T lymphocytes and thereby was ideal for investigating the role of NK cells in tumor. We analyzed the tissue-infiltrating NK cells in tumor and spleen using flow cytometry and bulk RNA sequencing (RNA-seq) (Figure S1A). Remarkably, compared to splenic NK cells, tumor-infiltrating NK cells displayed markedly reduced cytotoxicity (Figure 1A), effector function (Figure 1B and C), and polyfunction (Figure 1D). To further explore and compare the transcriptomic profiles of the intratumoral and splenic NK cells, we performed bulk RNA-seq of sorted tissue-infiltrating NK cells (Figure S1A). Principal component analysis (PCA) revealed the distinct transcriptional signatures between the tumor-infiltrating and splenic NK cells (Figure S1B). We observed a significant downregulation of 1734 genes in intratumoral NK cells compared to splenic NK cells, particularly those associated with NK cell activation and effector function (*Nkg7, Klrc1, Klrc2, Il12rb2, Slamf6, Il18r1, Pglyrp1*)^39,40^ (Figure 1E). Furthermore, gene set enrichment analysis (GSEA) revealed a considerably diminished enrichment of pathways and gene signatures associated with NK cell cytotoxicity and cytokines secretion in intratumoral NK cells (Figure 1F-I and Figure S3C). Altogether, these results provide compelling evidence of functional impairment in tumor-infiltrating NK cells compared to their counterparts in lymphoid organs.

**Figure 1.**
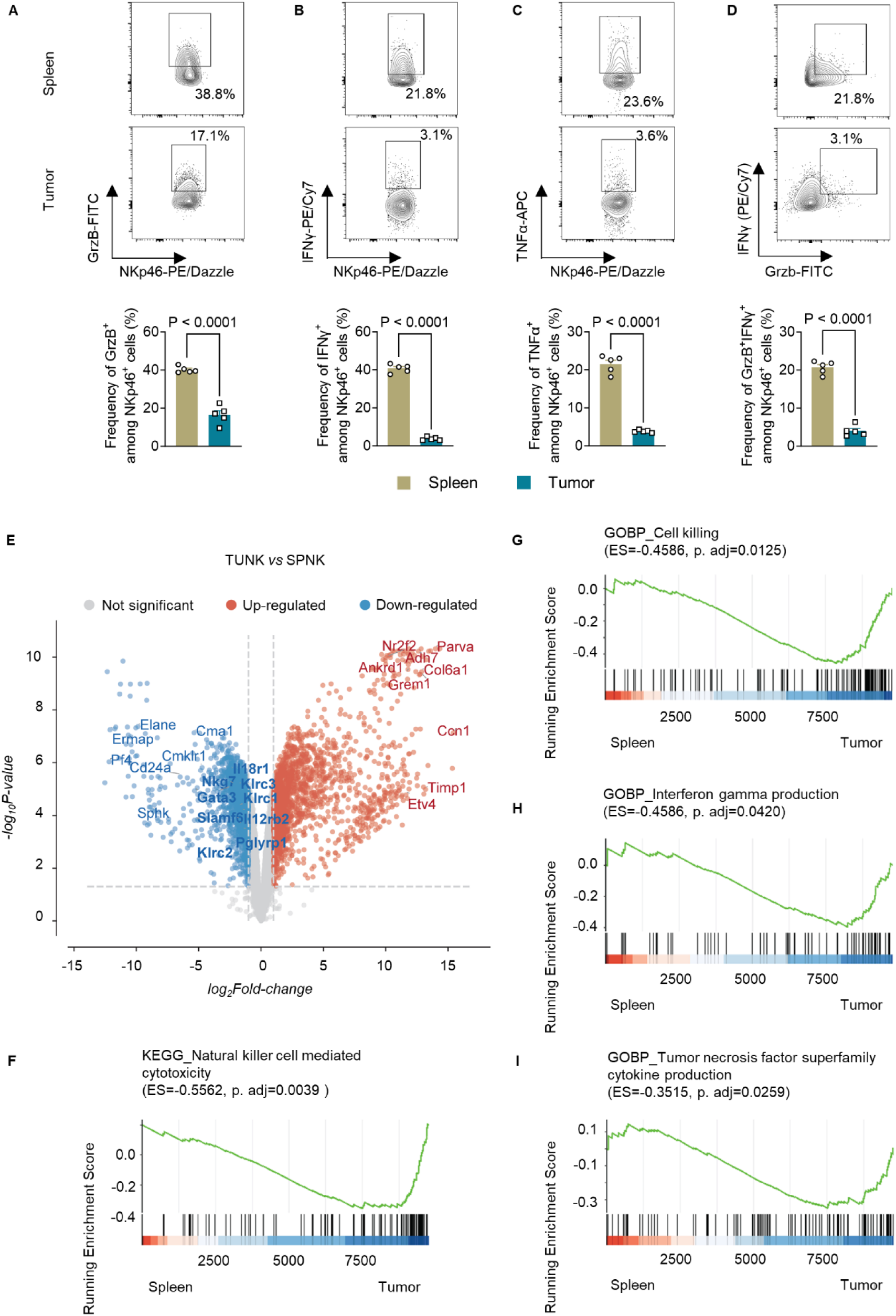
Tumor-infiltrating NK cells are functionally impaired. (A-D) BALB/c mice were inoculated with CT26_β2m^-/-^ tumor (8 × 10^5^, s.c.), and sacrificed two weeks post tumor inoculation, followed by flow cytometry analysis of NK cells (defined as CD3^-^NKp46^+^) in the spleen and tumor (n = 5 animals). Also see Figure S1A. Shown are representative flow cytometry plots and frequencies of GrzB^+^ (A), IFNγ^+^ (B), TNFα^+^ (C), and GrzB^+^IFNγ^+^ polyfunctional NK cells (D) among the total NK cells in spleen and tumor. Data represent the mean ± s.e.m. and are analyzed by two-sided Student’s t-test. (E-I) The experimental setting was similar to that shown in Fig. 1A-D except that NK cells were sorted from spleen (SPNK) or tumor (TUNK) for profiling gene expression with bulk RNA-seq (n = 4 animals). Pathway analysis was performed via GSEA with cluster profiler and mSigDB using KEGG and GO Biological Process Pathways. (E) Volcano plots representing upregulated and downregulated transcripts of TUNK *vs.* SPNK. Shown are the fold change in gene expression (log2(TUNK/SPNK)). The size of each data point is calculated based on −log10(p-value) × log2(FC), with a p-value cutoff of p < 0.05 and fold change cutoff of > −2 or < 2. Red dots indicate upregulated transcripts, blue dots represent downregulated transcripts, and bold-highlighted genes indicate those of interest. (F-I) GSEA plots of natural killer cell-mediated cytotoxicity (F), cell killing (G), interferon gamma production (H), and tumor necrosis factor superfamily cytokine production (I).

### IL-21−Fc reinvigorates exhausted NK cells for enhanced antitumor efficacy

To investigate the potential of IL-21 in rejuvenating exhausted NK cells within tumors, we first generated a recombinant protein by fusing murine IL-21 with a mutant mouse IgG2a (IL-21−Fc)^41-43^. The fusion protein retained comparable bioactivity to native IL-21 but had an extended half-life *in vivo* (Figure S2A-D). In the presence of IL-21−Fc in a co-culture experiment, NK cells exhibited enhanced killing efficiency toward various MHC-I-deficient tumor cells including CT26_β2m^-/-^ murine colorectal carcinoma cells, B16F10_β2m^-/-^ melanoma cells, and RMA-S lymphoma cells^44,45^. We noticed that *in vitro* treatment with IL-21−Fc markedly increased the expression of CD107a, as well as the production of granzyme B (GrzB) and interferon γ (IFNγ) (Figure 2A-D). Next, we investigated the effects of IL-21−Fc on tumor-infiltrating NK cells *in vivo*. Mice bearing CT26_β2m^-/-^ tumors (s.c.) were treated with multiple doses of intratumoral (i.t.) injection of IL-21−Fc or PBS solution as a control (Figure 2E). Interestingly, IL-21−Fc treatment did not lead to NK cell expansion (Figure 2F) or increased proliferative capacity (Figure 2G). Moreover, it had minimal impact on the apoptosis of tumor-infiltrating NK cells (Figure 2H). Instead, IL-21−Fc markedly enhanced the cytotoxicity and effector function of the intratumoral NK cells evidenced by the increased expression of GrzB and IFNγ (Figure 2I and J). To explore whether other immune cells in the TME also responded to IL-21−Fc, we examined immune cell infiltrates in the TME using flow cytometry. Notably, IL-21−Fc treatment showed minimum influence on other immune cell subsets except NK cells (Figure S2E). Collectively, our *in vitro* and *in vivo* results suggest that IL-21−Fc might revitalize the exhausted NK cells to enhance their effector function.

**Figure 2.**
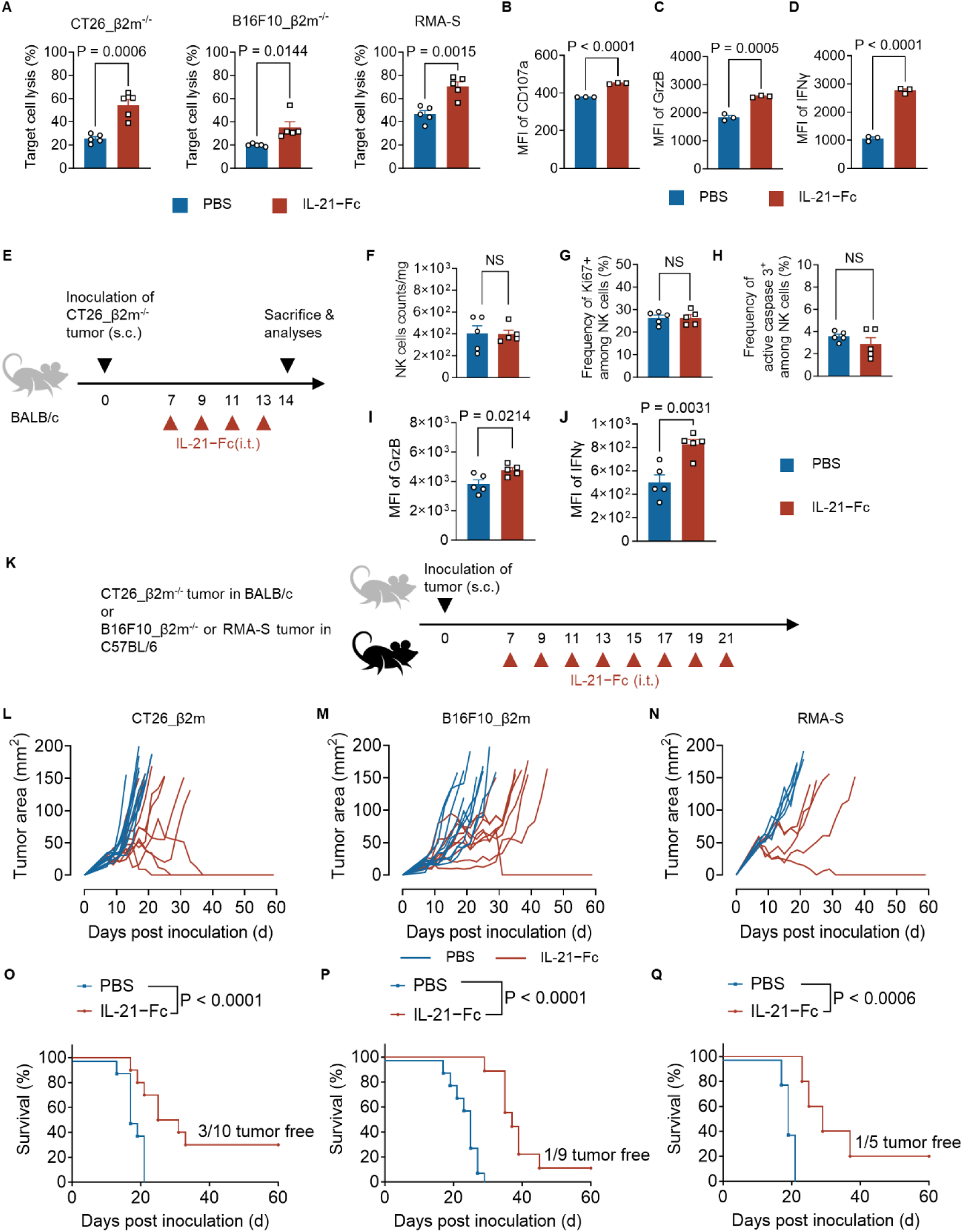
IL-21−Fc reinvigorates exhausted NK cells for enhanced antitumor efficacy. (A) Activated NK cells were co-cultured with indicated cancer cells at an effector-to-target (E/T) ratio of 0.5, in the presence or absence of IL-21−Fc (100 ng/mL) for 5 h. Shown is the percentage of lysis of target cells. (B-D) Activated NK cells were cultured in the presence or absence of IL-21−Fc (100 ng/mL) for 24 h, shown are the mean fluorescence intensity (MFI) of CD107a (**B**), GrzB (**C**), and IFNγ (**D**) of activated NK cells. (E) Experiment timeline for Fig. 2E-J. BALB/c mice were inoculated with CT26_β2m^-/-^ tumor cells (8 × 10^5^, s.c.) and received administration of IL-21−Fc (20 µg, i.t.) every two days for 4 doses in total. Mice were sacrificed two weeks post tumor inoculation, followed by flow cytometry analysis of NK cells in the tumor (n = 5 animals). (F) Counts of NK cells in tumors. (G-H) Frequency of Ki67^+^ (G) and active caspase 3^+^ (H) NK cells among tumor-infiltrating NK cells. (I-J) MFI of GrzB (I) and IFNγ (J) of tumor-infiltrating NK cells. (K) Experiment timeline for Fig. 2K-Q. BALB/c mice were inoculated with CT26_β2m^-/-^ colorectal carcinoma cells (5 × 10^5^, s.c.), and C57BL/6 mice were inoculated with B16F10_β2m^-/-^ melanoma cells (5 × 10^5^, s.c.), or RMA-S lymphoma cells (5 × 10^5^, s.c.), followed by administration of IL-21−Fc (20 µg, i.t.) or PBS for 8 doses in total (n = 5-10 animals). (L-N) Individual tumor growth curves of CT26_β2m^-/-^ colon carcinoma (L), B16F10_β2m^-/-^ melanoma (M), and RMA-S lymphoma (N). (O-Q) Kaplan-Meier survival curves of CT26_β2m^-/-^ colon carcinoma (O), B16F10_β2m^-/-^ melanoma (P), and RMA-S lymphoma (Q). All data represent the mean ± s.e.m. and are analyzed by two-sided Student’s t-test (A-D, F-J), or Log-rank test for survival curves (O-Q). NS, not significant (P > 0.05).

Encouraged by those results, we proceeded to evaluate the antitumor efficacy of IL-21−Fc in various syngeneic mouse tumor models (Figure 2K). We used immune-competent BALB/c or C57BL/6 mice to establish MHC-I-deficient s.c. tumor models including CT26_β2m^-/-^ colorectal carcinoma, B16F10_ β2m^-/-^ melanoma, and RMA-S lymphoma models, which did not respond to cytotoxic T lymphocyte-mediated killing, provided an ideal platform for accessing NK cell-mediated antitumor efficacy. Upon multiple injections of IL-21−Fc (i.t.), the cytokine monotherapy induced considerable tumor regression in treated mice, accompanied by prolonged survival (Figure 2L-N), thus demonstrating potent antitumor efficacy. Importantly, across different tumor models, 10-30 % of treated mice exhibited complete responses with established tumors fully eradicated (Figure 2O-Q). Therefore, IL-21−Fc emerges as an effective cytokine therapy capable of boosting NK cells *in vivo* to enhances antitumor efficacy against cancer cells that resist T cell-based immunotherapy.

### IL-21−Fc reprograms NK cell metabolism by elevating glycolysis

It has been reported that feeder cells expressing membrane-bound IL-21 increase Warburg metabolism of NK cells^36,46^. To investigate the underlying mechanism by which IL-21−Fc reinvigorates exhausted NK cells, we next investigated whether IL-21−Fc modulates NK cell metabolism. Activated NK cells cultured in the presence of IL-21−Fc exhibited greatly elevated basal and maximum extracellular acidification rate (ECAR) (Figure 3A-C), suggesting that IL-21−Fc promoted glycolytic activity of NK cells. Furthermore, IL-21−Fc treatment markedly increased glucose uptake, expression levels of the glucose transporter Glut1, and extracellular lactate concentration, providing additional evidence of enhanced NK cell glycolysis (Figure 3D-E and Figure S3A). Interestingly, the basal or maximal levels of oxygen consumption rates (OCR) of NK cells, which represent the OXPHOS activity, slightly decreased upon IL-21−Fc treatment (Figure S3B-D). Consequently, the ratio of ECAR to OCR was considerably increased, suggesting that IL-21−Fc reprograms NK cell metabolism toward a more glycolysis-dependent state (Figure 3F). We next applied 2-Deoxy-D-glucose (2-DG), a general inhibitor for glycolysis, to examine the role of metabolic reprogramming in reinvigorating NK cells. Importantly, we found that blockade of glycolysis using 2-DG, not OXPHOS by oligomycin, abrogated IL-21−Fc’s effects on enhancing NK cell effector function and cytotoxicity (Figure 3G and H). These results suggest that enhancement of glycolysis is critical for rejuvenating NK cells by IL-21−Fc.

**Figure 3.**
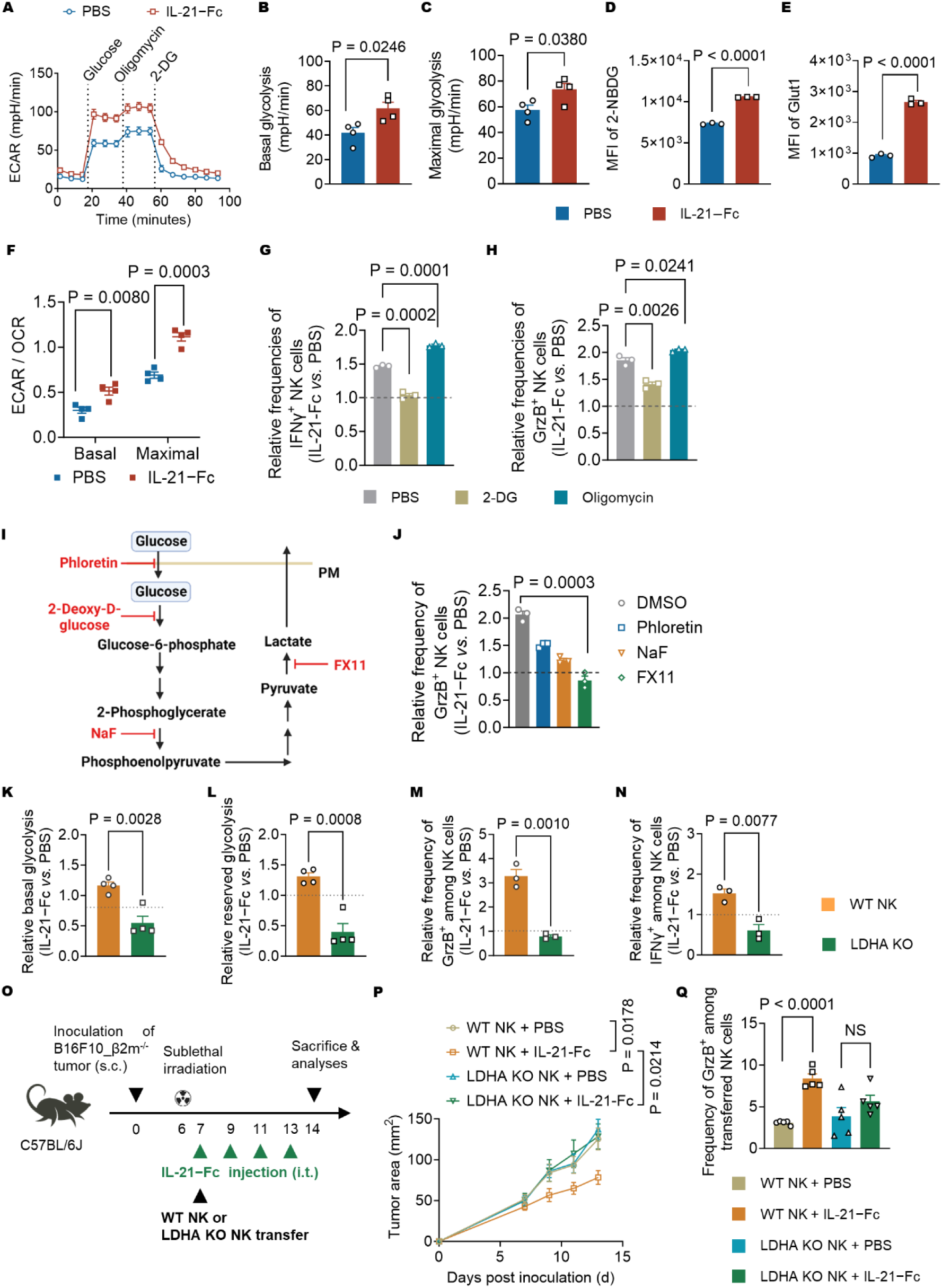
IL-21−Fc reprograms NK cell metabolism by elevating LDHA-dependent glycolysis. (A-F) Activated NK cells were cultured in the presence or absence of IL-21−Fc (100 ng/mL) for 24 h. (A) The real-time ECAR curves (n = 4 replicates). (B) The basal ECAR of NK cells. (C) The maximum ECAR of NK cells. (D) The MFI of 2-NBDG of NK cells. (E) The MFI of Glut1 of NK cells. (F) The ratio of basal and maximal ECAR to OCR of NK cells. (G-H) Activated NK cells were cultured in the presence or absence of IL-21−Fc (100 ng/mL) for 24 h, with supplementation of metabolic inhibitors. 2-DG, 0.5 mM; oligomycin, 1 µM. Shown are the relative frequency of IFNγ^+^ (G) and GrzB^+^ (H) NK cells among total NK cells in the IL-21−Fc treatment group normalized by that in the PBS group. (I) Schematic illustration of related metabolic pathways in glycolysis and inhibitors used. (J) Experiment setting was similar as that described in Fig. 3G-H. Phloretin, 50 μM; NaF, 4 mM; FX11, 20 μM. Shown are the relative frequency of GrzB^+^ NK cells among total NK cells in the IL-21−Fc treatment group normalized by that in the PBS group. (K-N) Generation of LDHA KO NK cells, also see Figure S3E. Shown are the relative basal glycolysis (K), relative reserved glycolysis (L), relative frequency of GrzB^+^ NK cells (M), and relative frequency of IFNγ^+^ NK cells (N) among total NK cells in the IL-21−Fc treatment group normalized by that in the PBS group. (O) The experiment timeline for Fig. 3O-Q. CD45.2^+^ C57BL/6J mice were inoculated with B16F10_β2m^-/-^ tumor cells (8 × 10^5^, s.c.), followed by sublethal irradiation on day 6 and adoptive transfer of Thy1.1^+^ WT NK cells (1 × 10^6^) or Thy1.1^+^ LDHA KO NK cells (1 × 10^6^) on day 7. Mice received injection of IL-21−Fc (20 µg, i.t.) or PBS as control every two days from day 7 for 4 doses in total. On day 14, mice were sacrificed followed by flow cytometry analysis of NK cells in tumors (n = 5 animals). (P) Average tumor growth curves of mice. (Q) The frequency of GrzB^+^ NK cells among transferred NK cells. All data represent the mean ± s.e.m. and are analyzed by two-sided Student’s t-test (A-H, J-N) or one-way ANOVA with Tukey’s test (P-Q).

To probe the molecular basis of IL-21−Fc-mediated NK cell reinvigoration, we employed specific inhibitors targeting factors involved in the glycolysis pathway and assessed NK cell responses (Figure 3I). We found that in the presence of an LDHA inhibitor, FX 11, IL-21−Fc-mediated enhancement of cytotoxicity of NK cells was completely abolished (Figure 3J). LDHA has been reported to play a central role in converting pyruvate to lactate for anaerobic metabolism, a process critical for NK cell function^46^. Subsequently, we generated LDHA knock-out (KO) NK cells using LDHA-targeting single-strand guide RNA (sgRNA) and the technology of Clustered Regularly Interspaced Short Palindromic Repeats (CRISPR)-Cas9 gene editing (Figure S3E and F). Compared to the wild-type (WT) NK cells (treated with scramble sgRNA), LDHA KO NK cells failed to respond to IL-21−Fc for enhanced glycolysis, cytotoxicity, or effector function (Figure 3K-N).

To further investigate the role of LDHA *in vivo*, we adoptively transferred WT or LDHA KO NK cells to mice bearing established B16F10_β2m^-/-^ tumors one day post lymphodepletion, followed by injection of IL-21−Fc or PBS (i.t.) (Figure 3O). While the combination therapy of transfer of WT NK cells and IL-21−Fc induced considerable tumor regression, LDHA KO NK cells in combination with IL-21−Fc failed to control the tumor growth (Figure 3P and Figure S3G). Flow cytometry analysis of tumor-infiltrating NK cells revealed that LDHA KO NK cells did not respond to the *in vivo* treatment of IL-21−Fc for increased cytotoxicity or effector function, thereby failing to eliminate MHC-I-deficient tumor cells (Figure 3Q and Figure 3H-J). Altogether, IL-21−Fc promotes LDHA-dependent glycolysis in exhausted NK cells, an effect that is indispensable for their reinvigoration with enhanced function and antitumor efficacy.

### IL-21−Fc combined with IL-15SA eradicates MHC-I-deficient tumors for durable protection

The observation that IL-21−Fc revitalizes intratumoral NK cells motivated us to assess the combination therapy of IL-21−Fc with IL-15SA, a widely used cytokine in NK cell-based therapy^47,48^. While IL-15 has been reported to effectively boost NK cell expansion, its use may lead to NK cell exhaustion and potentially life-threatening side effects in clinical settings^49,50^. To assess whether IL-21−Fc may synergize with IL-15SA in antitumor therapy, we co-administrated IL-21−Fc (20 µg × 8, i.t.) together with a low dosage of IL-15SA (5 µg × 2, i.t.) in several established MHC-I-deficient tumor models (Figure 4A). In the CT26_β2m^-/-^ tumor (s.c.) model, the combination therapy of IL-21−Fc and IL-15SA remarkably regressed tumor and induced durable cure in nearly 40 % of the treated mice, while IL-15SA alone failed to induce any complete responses (Figure 4B and E). Importantly, with the low dosage of IL-15SA, no overt systemic toxicity was observed during treatment, as evidenced by stable body weight in treated mice (Figure S4A). Intriguingly, most of the tumor-free mice from the combination therapy group rejected a rechallenge with CT26_β2m^-/-^ tumor cells (Figure S4D), suggesting that the cures mice might obtain memory-like antitumor immunity. These results together suggest the promise of IL-21−Fc and IL-15SA combination as a safe and effective therapy against a T cell-resistant tumor.

**Figure 4.**
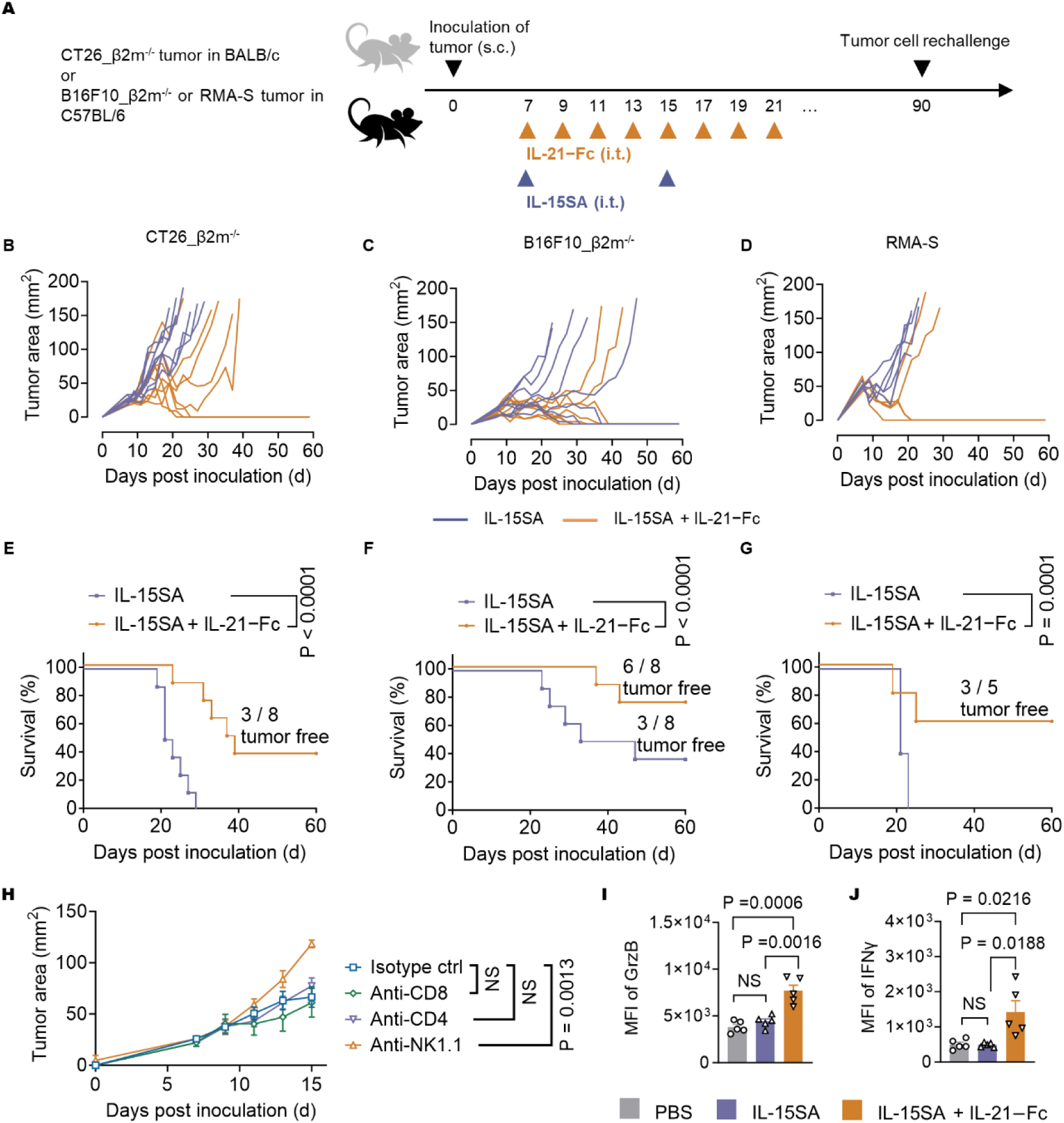
IL-21−Fc combined with IL-15SA eradicates MHC-I-deficient tumors and induces durable protection. (A) Experiment timeline. BALB/c mice were inoculated with CT26_β2m^-/-^ colon carcinoma cells (5 × 10^5^, s.c.), and C57BL/6J mice were inoculated with B16F10_β2m^-/-^ melanoma cells (5 × 10^5^, s.c.), or RMA-S lymphoma cells (8 × 10^5^, s.c.). Mice received administration of IL-15SA (5 µg, i.t.) on day 7 and day 14, and IL-21−Fc (20 µg, i.t.) or PBS every two days from day 7 for 8 doses in total. Cured mice from IL-15SA and IL-21−Fc combination groups were rechallenged with injection of CT26_β2m^-/-^ (1 × 10^5^, s.c.), or B16F10_β2m^-/-^ (1 × 10^5^, s.c.) cells 90 days post primary tumor inoculation. (B-D) Individual tumor growth curves of CT26_β2m^-/-^ colon carcinoma (B), B16F10_β2m^-/-^ melanoma (C), and RMA-S lymphoma (D). (E-G) Kaplan-Meier survival curves of CT26_β2m^-/-^ colon carcinoma (E), B16F10_β2m^-/-^ melanoma (F), and RMA-S lymphoma (G). (H) C57BL/6J mice were inoculated with B16F10_β2m^-/-^ melanoma cells (5 × 10^5^, s.c.), and received similar treatment as described in Figure 4A excepted for additional injection of immune cell depletion antibodies (400 µg, i.p.) every three days from day 5. Shown are the average tumor growth curves, also see Figure S4F. (I-J) BALB/c mice were inoculated with CT26_β2m^-/-^ colon carcinoma cells (5 × 10^5^, s.c.), followed by administration of IL-15SA (5 µg, i.t.) on day 7 and IL-21−Fc (20 µg) every two days from day 7 for 4 doses in total. On day 14, mice were sacrificed followed by flow cytometry analysis of NK cells in tumor. The control group receiving PBS administration was shared with the experiment described in Figure 2E-J following the 3R principle in animal welfare (n = 5 animals). Shown are the MFI of GrzB (I) and IFNγ (J) of tumor-infiltrating NK cells. All data represent the mean ± s.e.m. and are analyzed by Log-rank test for survival curves (E-G), or one-way ANOVA with Tukey’s test (H-J).

Extending the combinatory treatment to a highly aggressive B16F10_β2m^-/-^ melanoma model owing to its poor immunogenicity, the same combination therapy led to effective tumor regression and durable cures in up to 75 % of treated mice, whereas IL-15SA alone induced only moderate responses (Figure 4C and F). Impressively, 83.3 % of cured mice from the group of combination therapy rejected a second challenge of the B16F10_β2m^-/-^ tumor cells (Figure S4E), suggesting the formation of robust memory-like antitumor immunity. To further test the robustness of the combination therapy, we employed RMA-S cells to establish a lymphoma model (s.c.) and initiated the treatment when the tumors were established with big size (around 50 mm^2^). The combination of IL-21−Fc and IL-15SA resulted in a substantial regression in tumor growth and curative responses in 60 % of the treated mice, while IL-15SA monotherapy only transiently controlled tumor growth without complete eradication (Figure 4D and G). Notably, the combination therapy did not induce any systemic side effects in both B16F10_β2m^-/-^ and RMA-S models (Figure S4B and C).

To elucidate the immune cells contributing to the observed antitumor efficacy, we employed different antibodies to selectively deplete specific immune cell types (Figure S4F and G). Remarkably, only NK cell depletion using anit-NK1.1 antibody abolished the antitumor efficacy of combination treatment, while depletion of CD4^+^ T cells or CD8^+^ T cells had negligible effect (Figure 4H), providing evidence that NK cells are indispensable for the antitumor efficacy mediated by combination therapy of IL-21−Fc and IL-15SA. In addition, we found the combination therapy of IL-21−Fc and IL-15SA markedly promoted the cytotoxicity and effector function of tumor-infiltrating NK cells, whereas IL-15SA monotherapy showed negligible effects on NK cell rejuvenation in tumor (Figure 4I and J, Figure S4H). These results collectively imply that IL-21−Fc potentiates IL-15SA in NK cell-based therapy and demonstrate the promising antitumor activity mediated by IL-21−Fc through directly enhancing effector function of intratumoral NK cell *in vivo*.

### IL-21−Fc modulates NK cell transcription for enhanced activation, cytotoxicity, and effector function

To explore the role of IL-21−Fc in regulating intratumoral NK cells at the transcriptional level, we conducted single-cell RNA sequencing (scRNA-seq) analysis on NK cells (CD45.2^+^CD3^-^NKp46^+^) that were isolated from CT26_β2m^-/-^ tumors (Figure S5A). Mice treated with IL-15SA alone or in combination with IL-21−Fc were sacrificed, and intratumoral NK cells were sorted and pooled together within the same treatment. A total of 23,108 cells were analyzed (10,495 from the IL-15SA control group and 12,613 from the IL-15SA + IL-21−Fc combination group), with an average of 18,384 gene transcripts detected in the IL-15SA control group and 18,281 in the combination treated group. Unsupervised clustering unveiled four distinct clusters among tumor-infiltrating NK cells (Figure 5A and B). Comparative analysis revealed that, compared to control NK cells treated with IL-15SA alone, NK cells from mice receiving the combination treatment (IL-15SA + IL-21−Fc) exhibited significantly elevated expression of genes encoding activation and cytotoxic molecules, including *Ifitm, Spp1, Tnfrsf9, Gzmb,* and *Gzmc*^51^, along with glycolysis-related enzymes such as *Ldha* (Figure 5C). Remarkably, cluster 3 was predominantly enriched in the combination treatment group. This cluster displayed markedly heightened expression of genes encoding degranulation molecules, including *Gzmd, Gzmc,* and *Gzme* (Figure 5D and E, Figure S5B), and was strongly associated with glycolysis and cytotoxicity (Figure 5F and G). In addition, cluster 1 displayed a higher expression level of proliferative and cell cycle-associated genes, including *Tubb5, Pclaf, Top2a, and Tuba1b*, which was slightly enriched in the combination treatment group (Figure 5D and E, Figure S5B). Consistent with the observation that IL-21−Fc restored NK cell effector function by enhancing LDHA-dependent glycolysis, the scRNA-seq analysis revealed that combination treatment enriched population with upregulated glycolysis/gluconeogenesis, NK cell-mediated cytotoxicity (Figure 5H and I), as well as higher LDHA expression (Figure 5C). These findings support the notion that IL-21−Fc enhances glycolysis of intratumoral NK cells *in vivo* and promotes their cytotoxicity and effector function.

**Figure 5.**
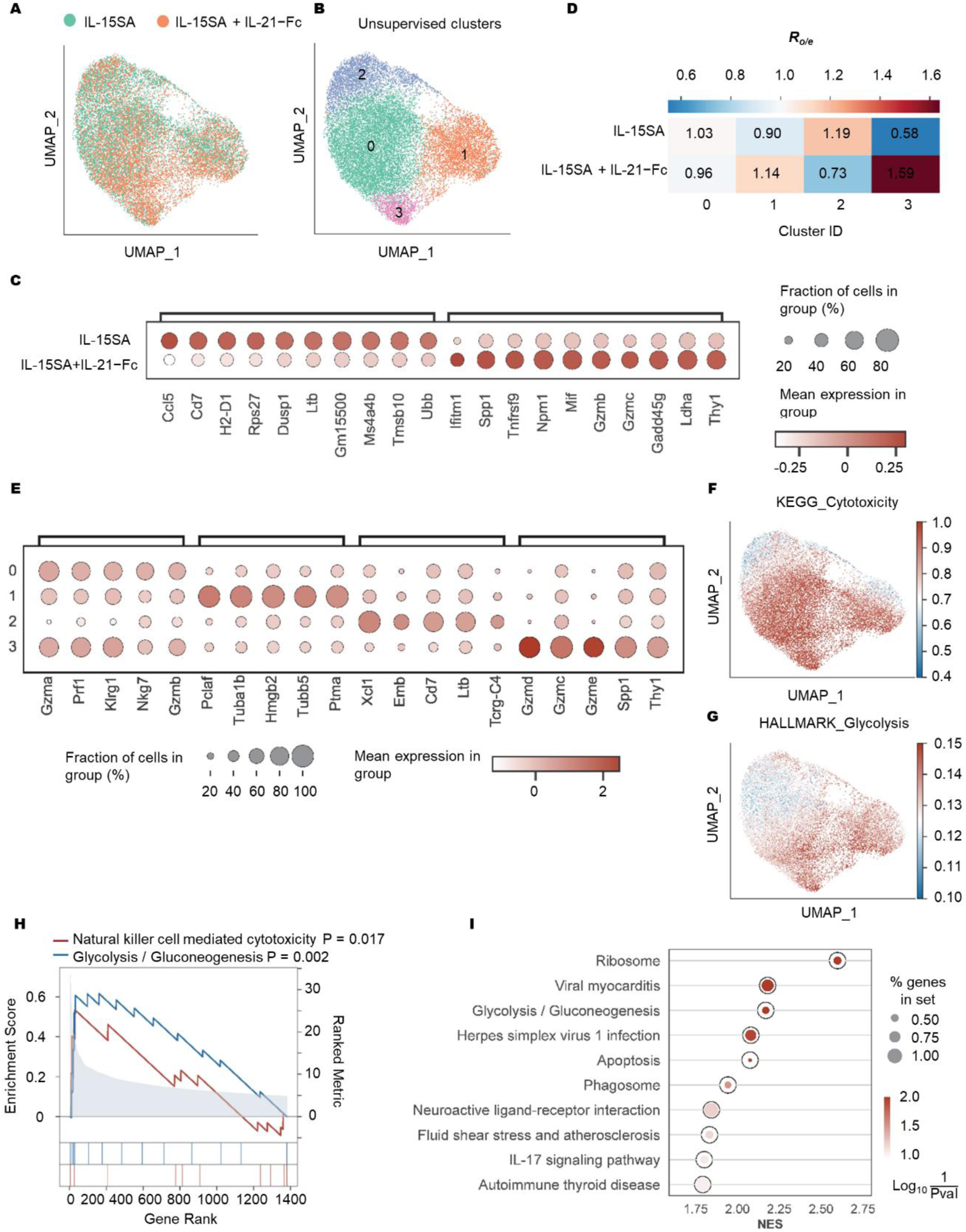
IL-21−Fc modulates NK cell transcription for enhanced activation, cytotoxicity, and effector function. (A-E) The experiment setting was similar as described in Figure 4I and J except that NK cells were sorted for scRNA-seq. Also see Figure S5A for experiment timeline. (A) Unsupervised UMAP clustering of NK cells sorted from CT26_β2m^-/-^ tumors in mice split by the treatment condition. (B) UMAP of four defining clusters of NK cells sorted from CT26_β2m^-/-^ tumors in mice. (C) The bubble plot representing expression features of the signature genes for treatment conditions. The fraction of cells in each group is indicated by the dot size and the mean expression levels of genes are indicated by the gradient color. (D) The ratio of observed and expected frequency of each cluster in Figure 5B. (E) The bubble plot representing expression features of the signature genes for each cluster. (F-G) UCell scoring of the KEGG-defining cytotoxicity pathway (F) and HALLMARK-defining glycolysis pathway (G). Scaled expression levels of the enrichment scores in each cluster are depicted using gradient color. (H) The multiple GSEA of NK cell-mediated cytotoxicity pathway and glycolysis/gluconeogenesis pathway between IL-15SA + IL-21−Fc and IL-15SA group of selected genes from the mSigDB database. (I) The signaling pathway regulated by direct comparison of differentially expressed genes (DEGs) between IL-15SA + IL-21−Fc and IL-15SA treatment. Pathway terms are ranked by –log10(p-value).

### IL-21−Fc enhances glycolysis of intratumoral NK cells *in vivo*

To further investigate the metabolic network alterations, we conducted unsupervised single-cell clustering analysis based on the pathway scores of 1075 genes involved in metabolism (Tabel S1). Compared to the NK cells from the control group receiving IL-15SA alone, NK cells from the combination treatment group exhibited enrichment in metabolic clusters 2, 3, and 4 (Figure 6A-C). In particular, genes encoding glycolysis/gluconeogenesis were significantly upregulated in cluster 3, which was enriched in the combination treatment group (Figure 6D).

**Figure 6.**
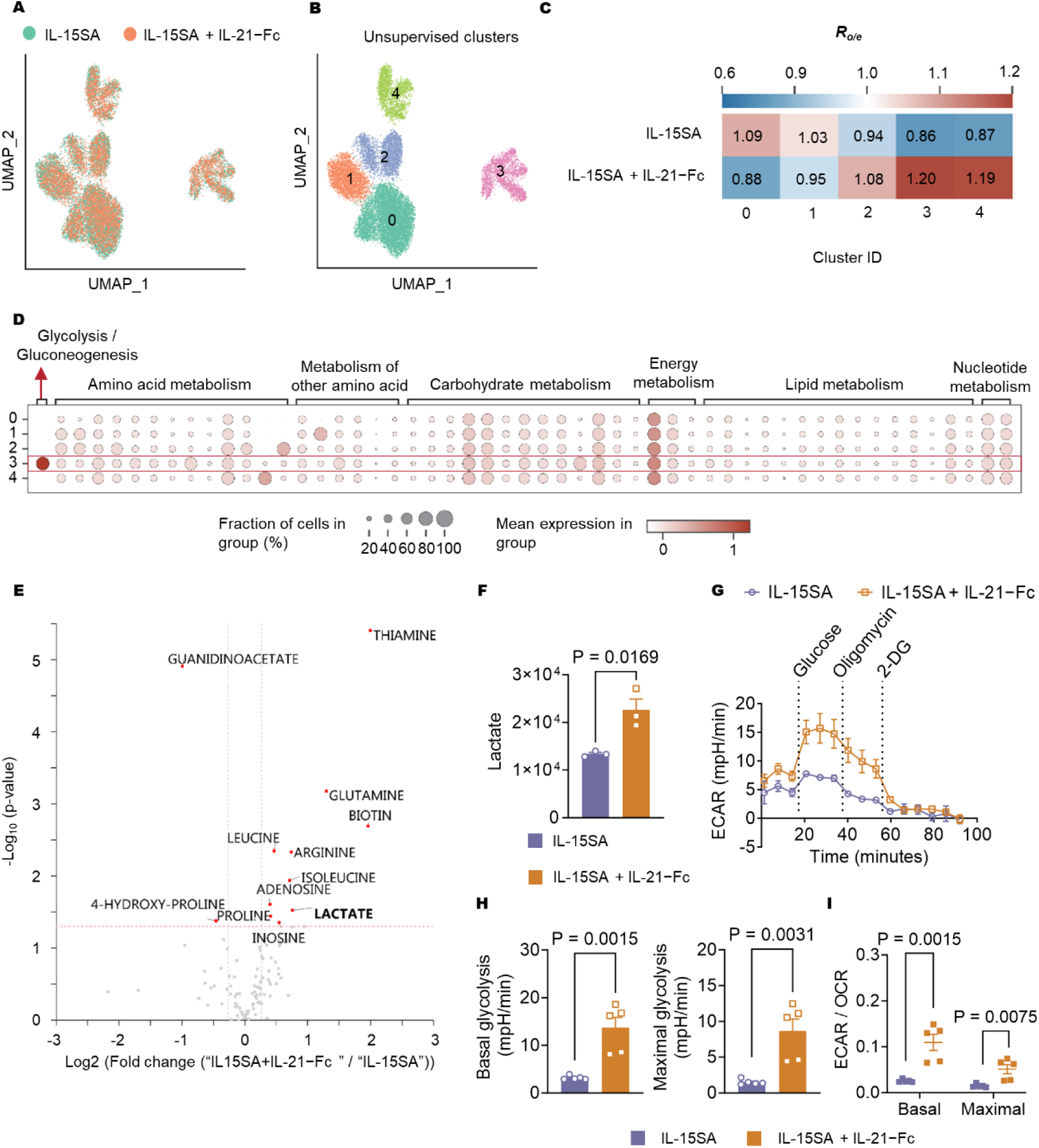
IL-21−Fc enhances glycolysis of intratumoral NK cells. (A) The unsupervised UMAP of NK cells sorted from tumors in mice split by the treatment condition based on the genes involved in KEGG-defining metabolic pathways. Also see Table S1. (B) UMAP of five defining clusters of NK cells sorted from tumors in mice based on the genes involved in KEGG-defining metabolic pathways. (C) The ratio of observed and expected frequency of each cluster in Fig. 6B. (D) The bubble plot representing expression features of the metabolic pathways for each cluster. (E) The volcano plot of top upregulated or downregulated metabolites among 110 detected metabolites of NK cells treated with IL-15SA + IL-21−Fc *vs.* IL-15SA, p-value cutoff p < 0.05. The experiment setting was similar to Figure 5 except that NK cells were sorted for metabolomics measurement (n = 3 animals). Also see Figure S6A. (F) The relative quantification of lactate based on extracted ion chromatogram. Also see Table S2. (G) The real-time ECAR curves. The experimental setting was similar to Fig. 5 except that NK cells were sorted for seahorse measurement (n = 3 animals). Also see Fig. S6A. (H) The basal and maximum ECAR of sorted NK cells. (I) The ratio of basal and maximal ECAR to OCR of sorted NK cells. All data represent the mean ± s.e.m. and are analyzed by two-sided Student’s t-test.

Subsequently, we conducted mass spectrometry-based metabolomics analysis to further study the metabolic reprogramming of NK cells mediated by IL-21−Fc *in vivo* (Figure S6A and B, Table S2). Several metabolites involved in glycolysis and amino acid metabolism, such as lactate, were markedly upregulated in intratumoral NK cells following combination therapy (Figure 6E and F). Moreover, the majority of metabolites associated with glycolysis showed noticeable increases in intratumoral NK cells upon combination treatment (Figure S6C).

In addition, we monitored the real-time changes of ECAR and OCR of intratumoral NK cells to assess the metabolic profiles *in vivo*. The tumor-infiltrating NK cells exhibited greatly enhanced basal and maximum ECAR upon the combination treatment (Figure 6G and H). Notably, the substantially increased ratio of ECAR to OCR indicated an IL-21−Fc-mediated metabolic shift towards glycolysis-dependent in these intratumoral NK cells (Figure 6I). Taken together, these results demonstrate that IL-21−Fc remarkably modulates the metabolic profiles of intratumoral NK cells, leading to enhanced glycolysis.

### IL-21−Fc reinvigorates human NK cells for enhanced antitumor efficacy in a xenograft model

Exhaustion-associated dysfunction of NK cells in human malignancies poses a significant challenge to the efficacy of NK-based immunotherapy^52-54^. Next, we aim to investigate whether IL-21 could also reinvigorate exhausted human NK (hNK) cells *in vivo* and promote their activity against xenografted tumors. We generated a fusion protein of human IL-21 (hIL-21) and human IgG1 Fc (hIL-21−Fc)^55,56^ (Figure S7A and B). hIL-21−Fc remarkably enhanced the activation and cytotoxic function of hNK cells derived from peripheral blood mononuclear cells (PBMC) of healthy donors. *In vitro* treatment with hIL-21−Fc led to increased expression of NKG2A, NKG2C, CD107a, Perforin and IFNγ in hNK cells (Figure 7A-E). Moreover, hNK exhibited enhanced target cell lysis against various human cancer cells, including MDA-MB-231 breast cancer cells, and U87 glioblastoma cells (Figure S7C and D). Consistent with the metabolic reprogramming effects of IL-21−Fc observed in mouse NK cells, hIL-21−Fc markedly elevated basal and maximum ECAR of hNK cells, as well as the ratio of ECAR to OCR (Figure 7F-I), whereas the OCR levels remained unchanged (Figure S7E-G).

**Figure 7.**
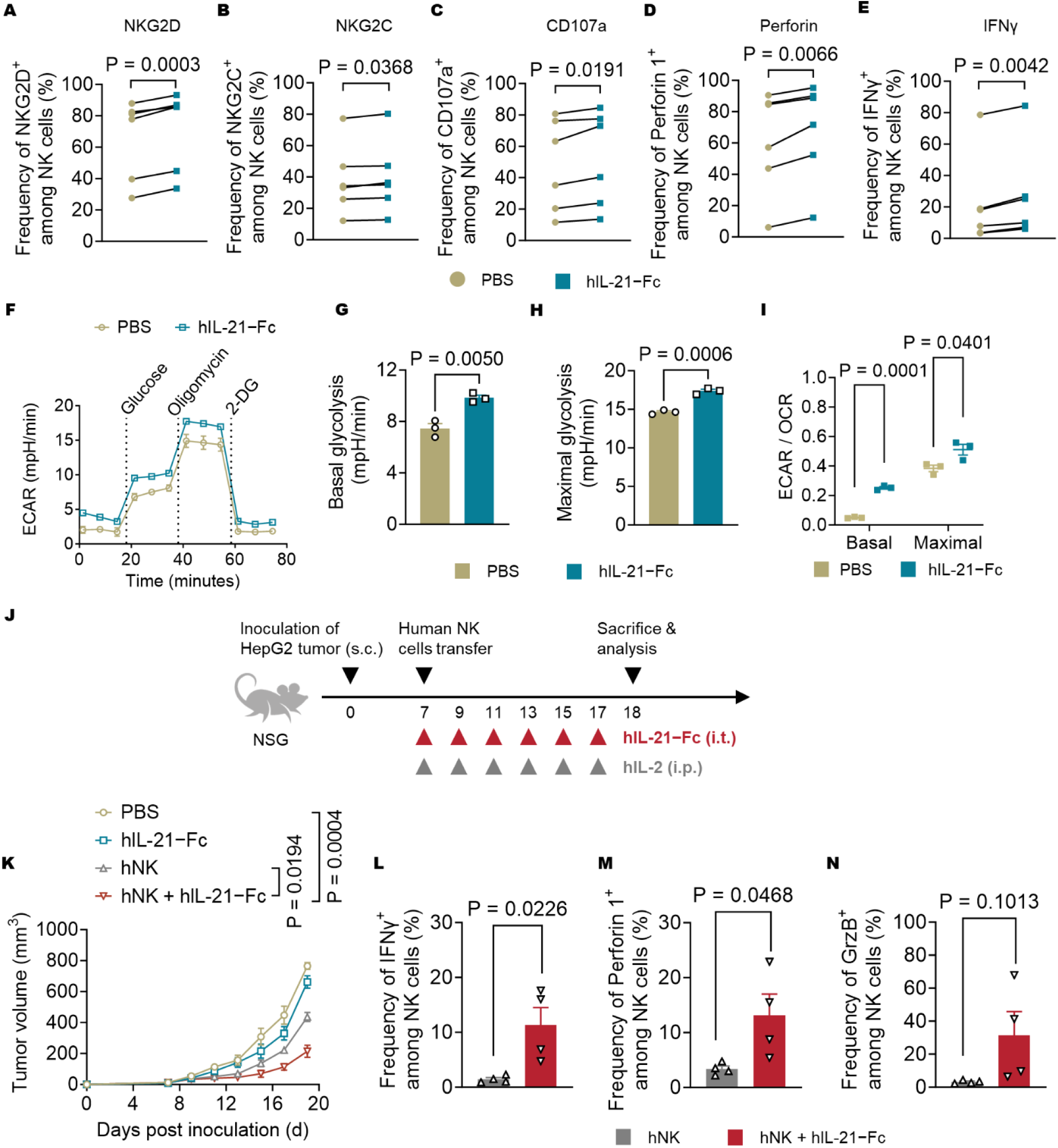
IL-21−Fc reinvigorates human NK cells for enhanced antitumor efficacy in a xenograft model. (A-E) hNK cells were isolated from the PBMC of healthy donors. PBMC-derived NK cells were preactivated with human IL-2 (100 U/mL) and IL-15 (20 ng/mL) for 24 h, followed by culturing in the presence or absence of hIL-21−Fc (100 ng/mL) (n = 3–6 donors). Shown are the frequency of NKG2D^+^ (A), NKG2C^+^ (B), CD107a^+^ (C), Perforin 1^+^ (D), and IFNγ^+^ (E) among total NK cells. (F) The real-time ECAR curves. (G) The basal ECAR of hNK cells. (H) The maximum ECAR of hNK cells. (I) The ratio of basal and maximal ECAR to OCR of hNK cells. (J) The experiment timeline. NSG mice were inoculated with HepG2 tumors (8 × 10^6^, s.c.), followed by transfer of hNK cells on day 7. Mice received administration of hIL-2 (75 kU, i.p.) and administration of hIL-21−Fc (20 µg, i.t.) every two days for 6 doses in total. Mice were sacrificed on day 18, followed by flow cytometry analysis of NK cells (defined as CD3^-^CD56^+^ cells) in tumors (n = 5 donors). (K) Average tumor growth curves. (L-N) Frequency of IFNγ^+^ (L), Perforin^+^ (M), and GrzB^+^ (N) NK cells among transferred NK cells. All data represent the mean ± s.e.m. and are analyzed by two-sided paired Student’s t-test (A-E), two-sided unpaired Student’s t-test (G-I, L-N), or one-way ANOVA with Tukey’s test (K).

We next assessed the antitumor activity of hIL-21−Fc in a xenografted HepG2 hepatocellular carcinoma model in NSG mice. Following inoculation of HepG2 tumor cells (5 × 10^6^, s.c.) in immunodeficient NSG mice, hNK cells were adoptively transferred 7 days post-tumor inoculation. Intraperitoneal (i.p.) injection of human interleukin-2 (hIL-2, 75 kU) was adopted to adjuvant the transferred NK cells^57^ (Figure 7J). Adoptive transfer of hNK cells alone exhibited limited efficacy in controlling tumor growth (Figure 7K). By contrast, mice received adoptive transfer of hNK cells in combination with administration of hIL-21−Fc (20 µg × 6, i.t.) showed noticeable tumor regression (Figure 7K), implying that hIL-21−Fc improved hNK cell activity against xenograft tumors. Subsequent analysis of tumor-infiltrating hNK cells revealed that similar to mouse syngeneic models, treatment with hIL-21−Fc greatly enhanced the effector function and cytotoxicity of hNK cells in the xenograft tumor, evidenced by increased frequencies of IFNγ^+^, Perforin 1^+^, and GrzB^+^ hNK cells (Figure 7L-N). These results collectively suggested that hIL-21−Fc reinvigorates hNK cells and potentiates NK cell therapy in human malignancies.

## DISCUSSION

NK cells are promising therapy to recognize and eliminate MHC-I-deficient tumors. However, their efficacy is often compromised by the immunosuppressive TME, leading to immune evasion and tumor progression. In this study, we demonstrate that a half-life extended recombinant cytokine, IL-21−Fc, offers a safe and effective *in vivo* therapeutic intervention for reinvigorating the tumor-infiltrating NK cells. Treatment with IL-21−Fc enhanced the effector function of mouse and human NK cells in syngeneic and xenograft tumor models, respectively, by promoting LDHA-dependent glycolysis. These findings offer novel insights into metabolic reprogramming mediated by IL-21−Fc and its potential to rejuvenate exhausted NK cells, paving the way for enhanced NK cell-based immunotherapy against cancer in clinical practice.

Current research efforts have primarily focused on using cytokines such as IL-21 for *ex vivo* expansion of NK cells for therapeutic application^35,58^. Several ongoing clinical trials are evaluating the efficacy of adoptive transfer therapy involving NK cells expanded *ex vivo* using feeder cells engineered to express membrane-bound IL-21^59,60^. However, a notable limitation of this approach is the requirement for complete removal of feeder cells prior to NK cell transfer, a process that poses challenges in Good Manufacturing Practice (GMP) facility^61,62^. Moreover, feeder cell-expanded NK cells still exhibit functional exhaustion within the TME^36,52^. Direct *in vivo* reinvigoration of exhausted NK cells using IL-21−Fc may provide a complementary therapeutic strategy to existing NK cell-based immunotherapies, potentially enhancing patients’ response rate.

IL-15 has been extensively utilized in both preclinical and clinical studies for NK cell-based immunotherapy due to its superior effects on NK cell expansion. However, dose-limiting side effects have been observed in clinical studies^50,63^. In addition, overexposure of NK cells to IL-15 induces NK cell exhaustion, thereby impairing NK cell therapy^49,64^. As an alternative approach, we utilized a low dose of IL-15SA that allowed for sufficient NK cell expansion *in vivo* without causing overt toxicity. Combination therapy involving IL-21−Fc and low-dose IL-15SA led to remarkable tumor regression and curative response in various MHC-I-deficient tumor models. Notably, multiple doses of IL-21−Fc were well tolerated by the treated mice. Together, these findings provide novel insights into the rational design of combinatory cytokine therapies aiming at directly boosting NK cells *in vivo*. Additionally, other cytokines that have been shown to boost NK cell expansion and/or function, including IL-2, IL-12, and IL-18^15,65-67^, may warrant consideration for future combination therapies involving *in vivo* interventions.

Cellular metabolism plays a pivotal role in regulating the fate of immune cells^24-26,68^. Glycolysis, in particular, is a critical metabolic pathway that fuels NK cells during activation and facilitates their effector function. Recent studies have highlighted the importance of glycolysis in NK cells function^69^, while glycolytic inhibition decreased the cytolytic function and their interactions with target cells^70^. In addition, activation of HIF1-α in NK cells within the TME has been shown to drive glycolysis, thereby enhancing their effector function and antitumor performance^71,72^. Our findings shown here indicate that the augmented antitumor function of NK cells mediated by IL-21−Fc relies on LDHA-dependent glycolysis. LDHA plays a crucial role in catalyzing the conversion of pyruvate to lactate, with the concomitant conversion of NADH to NAD^+73^, which in turn recycles NADH to NAD^+^ and reinforces glycolysis flux, thereby favoring NK cell function^74,75^. Therefore, modulating NK cell metabolism offers a promising strategy for controlling their function and ultimately improving the therapeutic outcomes of NK cell-based immunotherapy.

## ACKNOWLEDGEMENTS

We thank Dr. D.H. Raulet (University of California, Berkeley) for providing CT26_β2m^-/-^ and B16F10_β2m^-/-^ cells; Dr. W. Held (University of Lausanne) for providing RMA-S cells; D.J. Irvine (Massachusetts Institute of Technology) for providing the plasmid of mutant Fc domain; J. Auwerx (EPFL) for providing access to a Seahorse XFe96 Analyzer. We acknowledge the Metabolomics Platform at University of Lausanne, Swiss Institute of Bioinformatics, and Center of PhenoGenomics, Flow Cytometry Core Facility, Protein Expression Core Facility, and Gene Expression Core Facility at EPFL for technical assistance. L.T. acknowledges the grant support from Swiss National Science Foundation (315230_204202, IZLCZ0_206035, CRSII5_205930), European Research Council under the ERC grant agreement MechanoIMM (805337), Swiss Cancer Research Foundation (KFS-4600-08-2018), Kristian Gerhard Jebsen Foundation, Anna Fuller Fund, Xtalpi Inc., and EPFL. C.S. was supported by the National Key R&D Program of China (2021YFC2300604, 2022YFA1303200), National Natural Science Foundation of China (#82022056, #8239445, #92169118), CAS Project for Young Scientists in Basic Research (#YSBR-068). A.K. was supported by the European Union’s Horizon 2020 research and innovation program under the Marie Skłodowska-Curie grant agreement No. 754354. M.G. was supported by the Chinese Scholarship Council (CSC, No. 201808320453)

## AUTHOR CONTRIBUTIONS

Y.W., C.S., Y.G., and L.T. conceived the study. Y.W., C.S., Y.G., and L.T. designed the experiments. Y.W., C.H., G.C., M.A., A.K., Y.Z., B.F., M.G., S.C., Z.Z., C.S., and Y.G. performed the experiments. Y.W., C.H., C.S., Y.G., and L.T. analyzed the data. Y.W. and L.T. wrote the manuscript. All authors edited the manuscript.

## STAR★METHODS

Detailed methods are provided in the online version of this paper and include the following:

### KEY RESOURSE TABLE

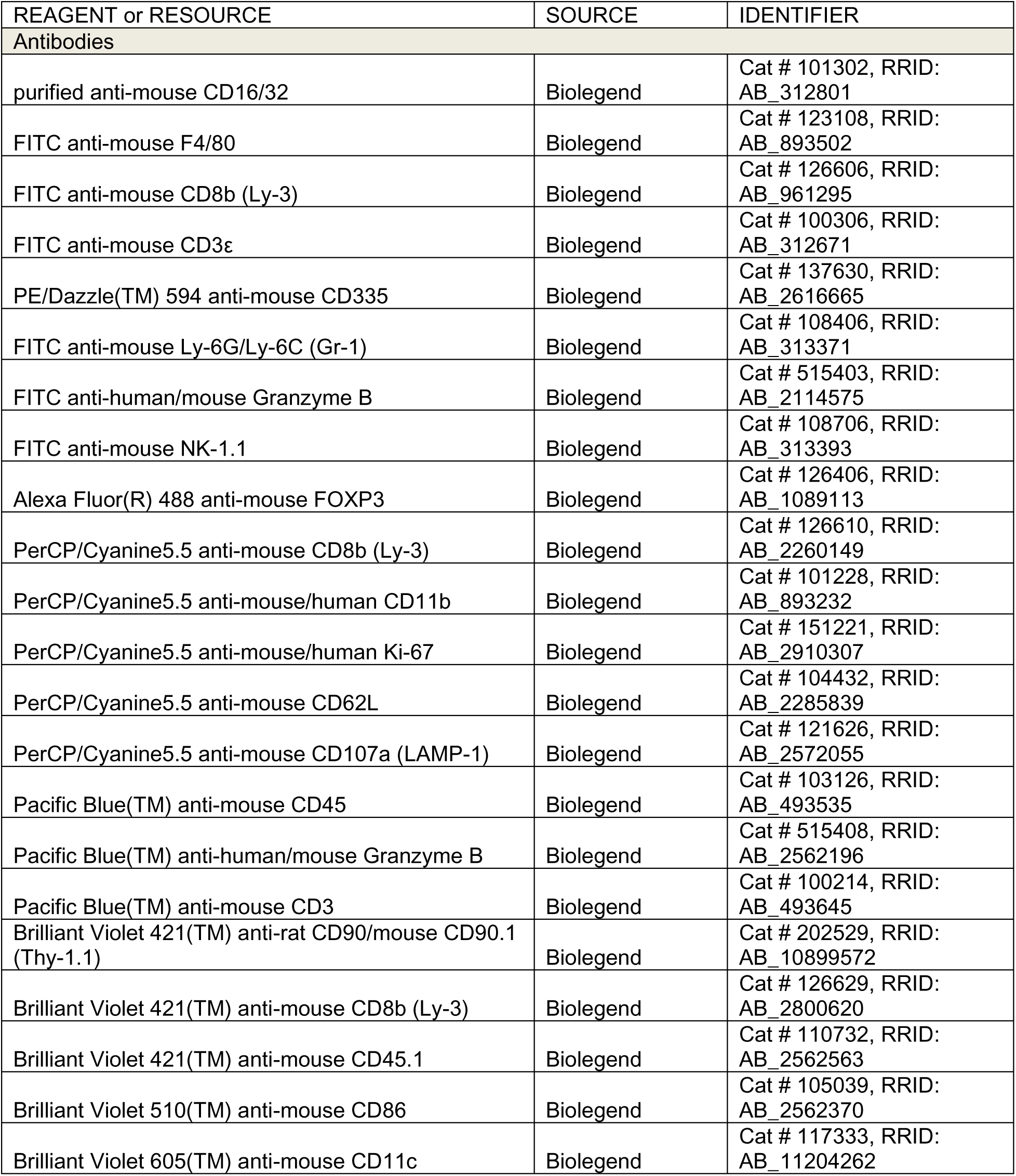
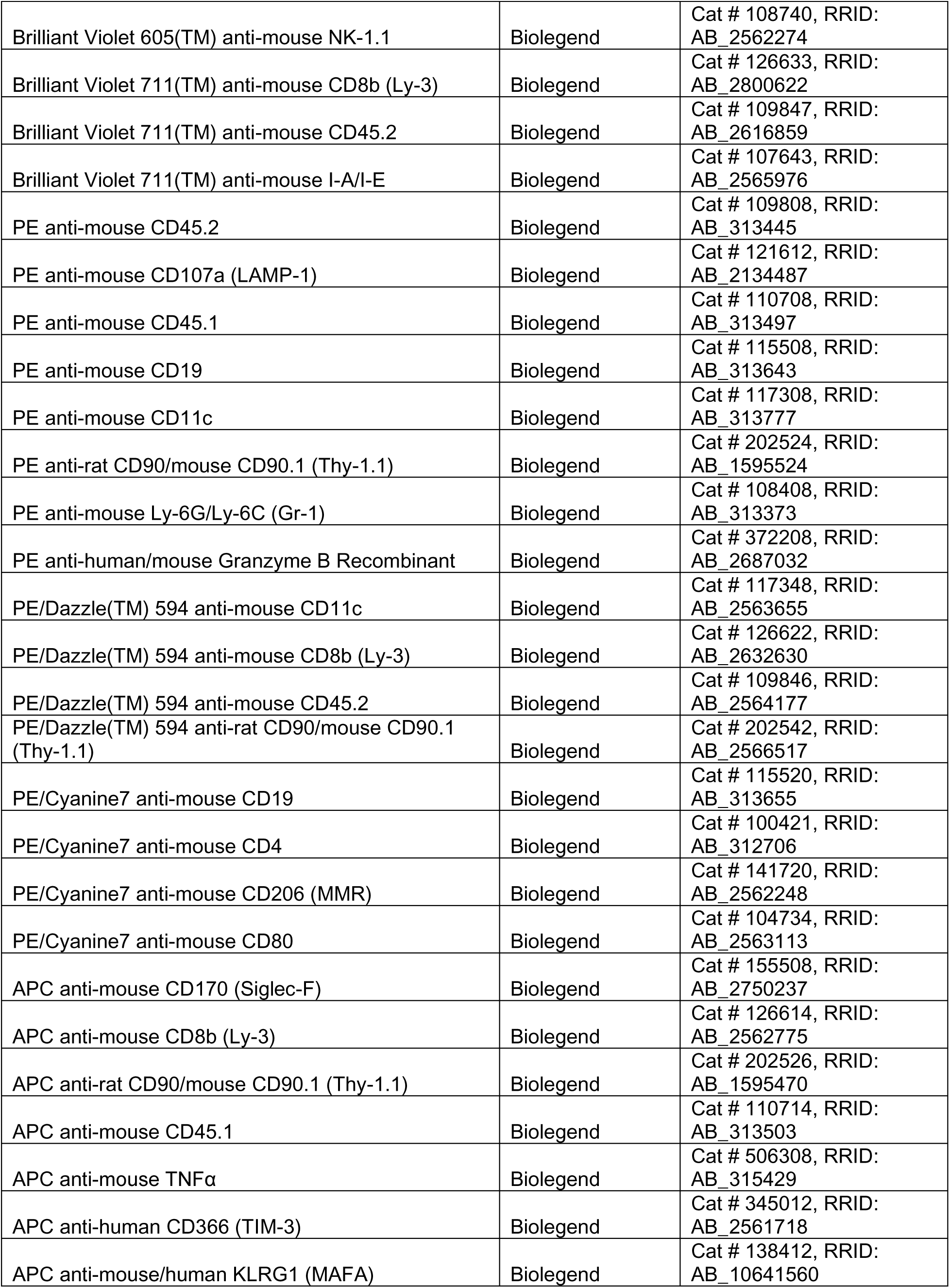
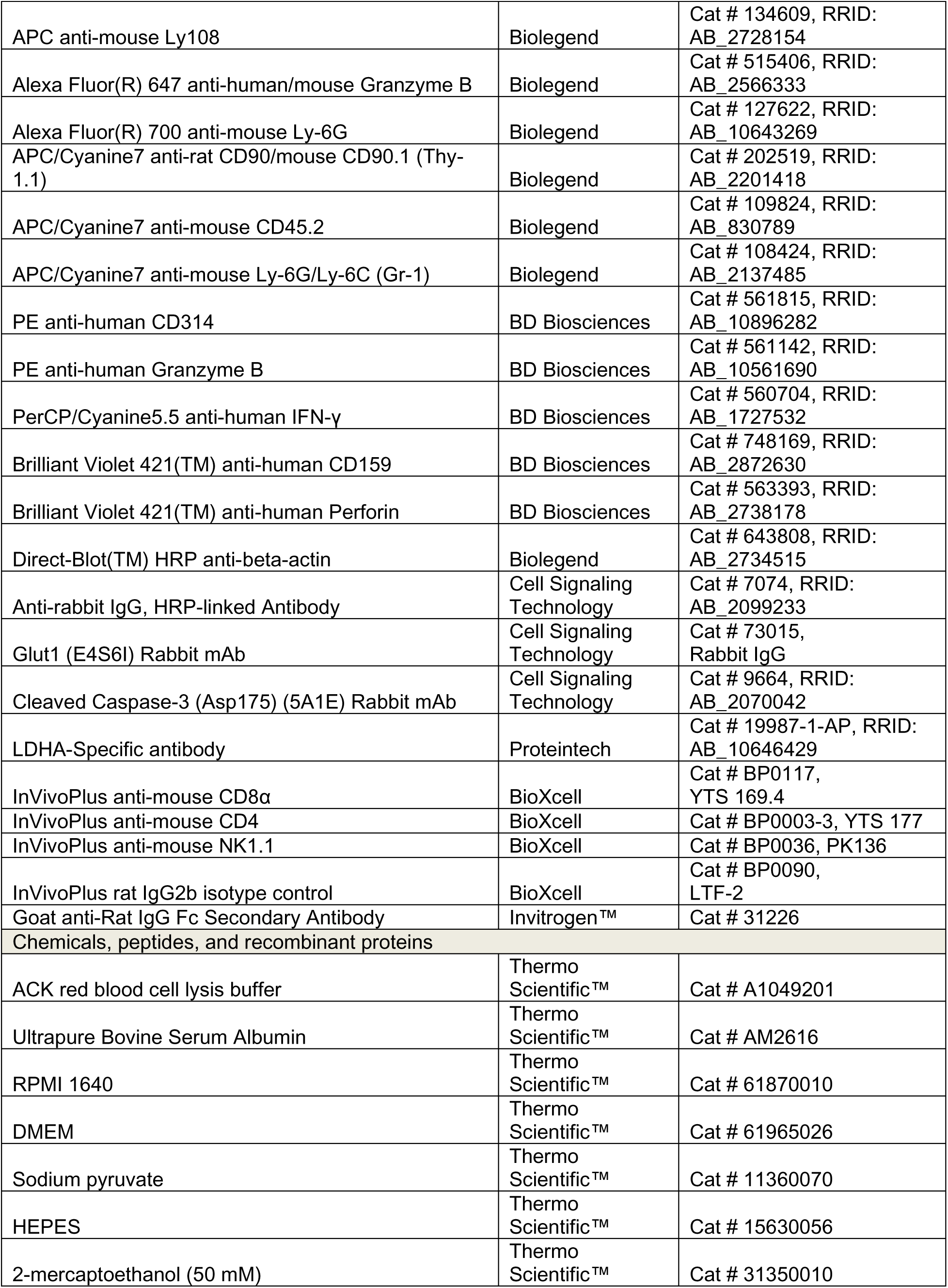
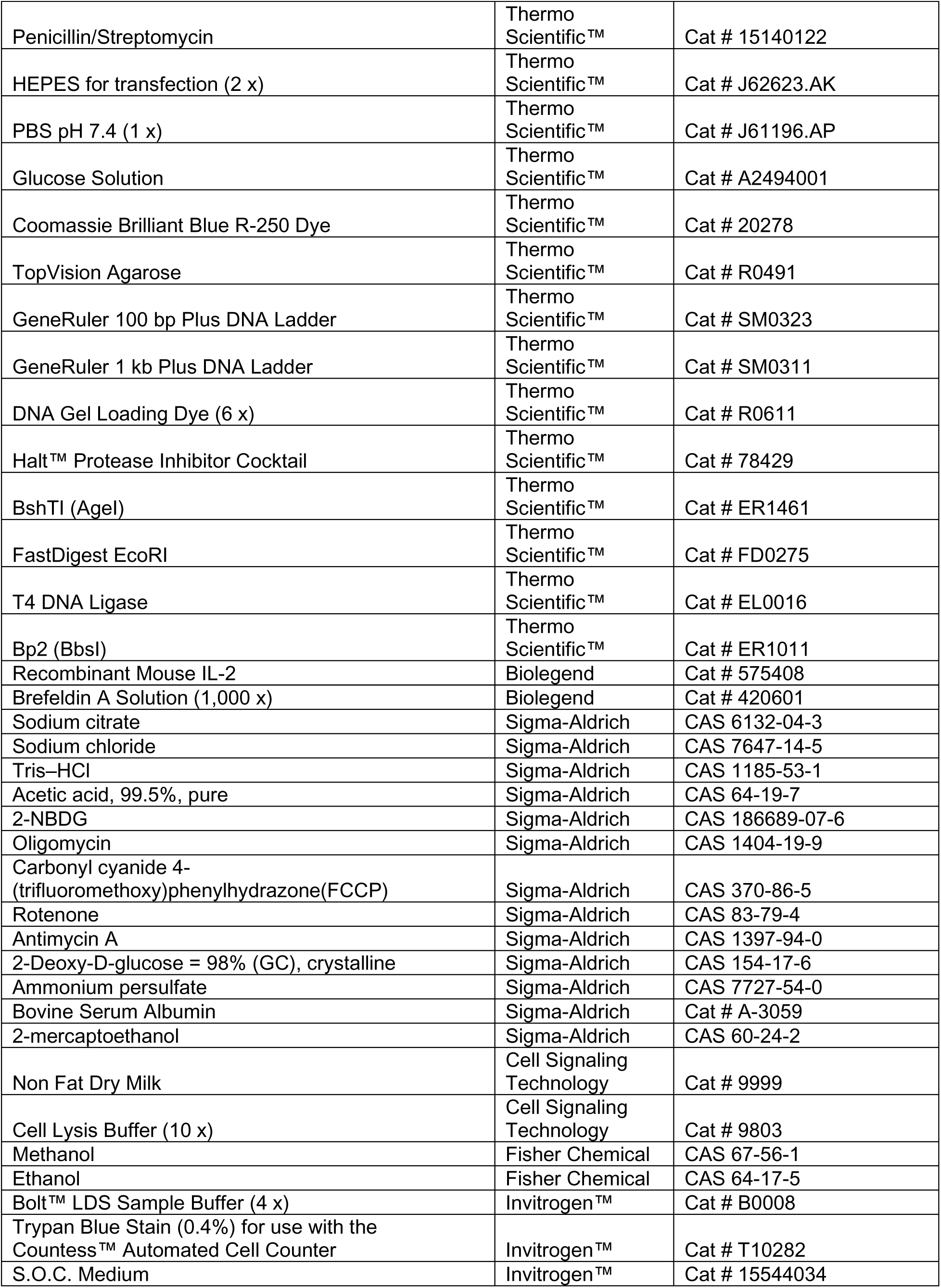
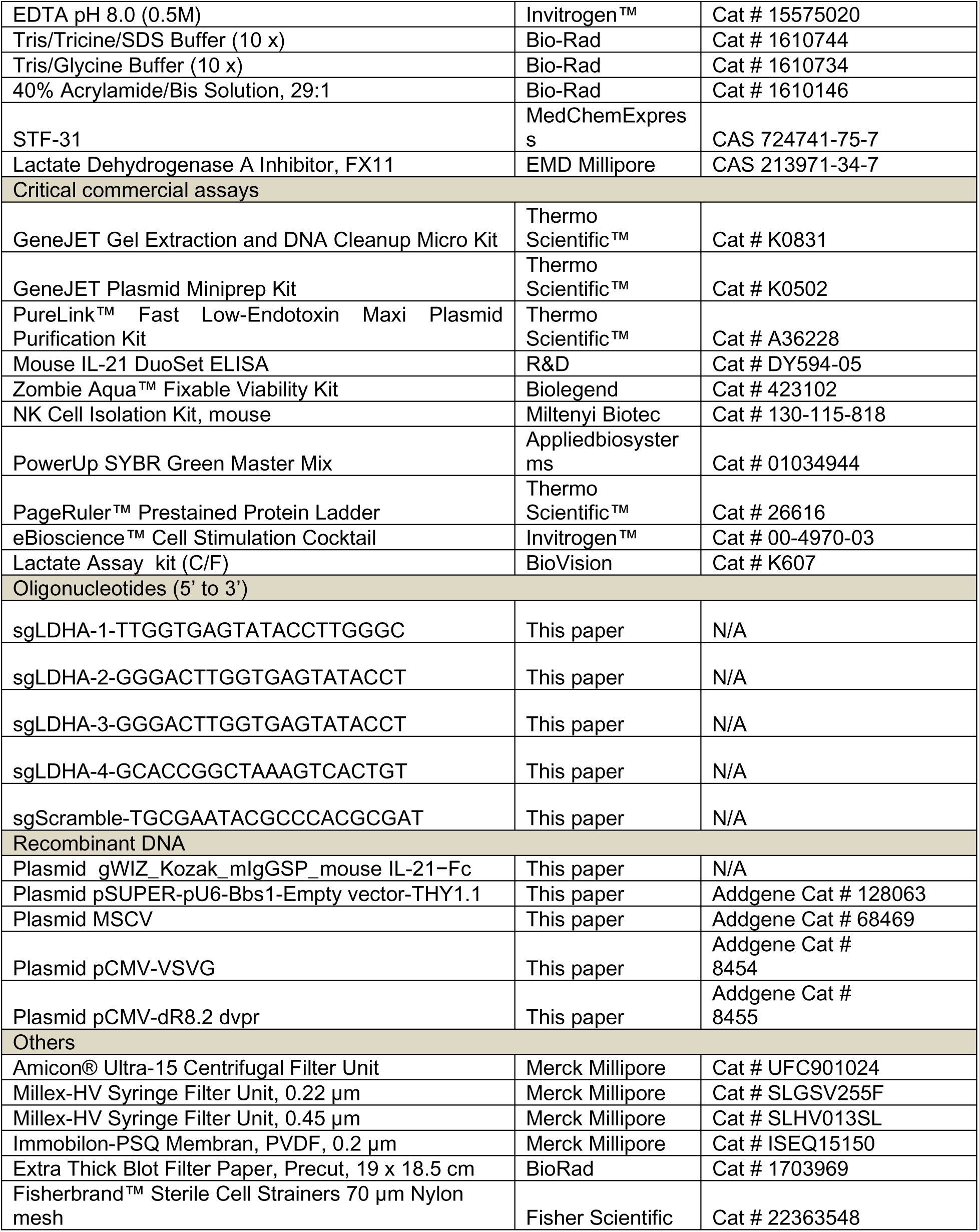

### RESOURSE AVAILABILITY

#### Lead contact

Further information and requests for resources and reagents should be directed to and will be fulfilled by the lead contact, Li Tang

#### Materials availability

Requests for materials should be directed to the lead contact, Li Tang. Material transfer agreements will be necessary to obtain reagents.

#### Data and code availability

The RNA-seq data for tumor-infiltrating lymphocytes are available in the Gene Expression Omnibus (GEO) database under accession number GSE xxx (to be updated when published). All relevant data are available from the corresponding author upon request. Accession numbers are listed in the key resource table. Any additional information required to the reanalyze the data reported in this paper is available from the lead contact upon request.

### EXPERIMENTAL MODEL AND STUDY PARTICIPANT DETAILS

#### Cell lines

B16F10_β2m^-/-^ and CT26_β2m^-/-^ cells were provided by Prof. David H. Raulet from the University of California, Berkeley. RMA-S cell was provided by Prof. Werner Held from the University of Lausanne. U87, MDA-MB-231, and Phoenix-Eco cells were originally acquired from the American Type Culture Collection (Manassas, VA, USA). HepG2 cells were originally purchased from Haixing Biosciences (Suzhou, Jiangsu, China). B16F10_β2m^-/-^, MDA-MB-231, and Phoenix-Eco cells were cultured in DEME medium (Thermo Fisher Scientific) supplemented with Fetal Bovine Serum (FBS) (10 % v/v, Thermo Fisher Scientific), HEPES (1 % v/v, Thermo Fisher Scientific), sodium pyruvate (1 % v/v, Thermo Fisher Scientific), penicillin/streptomycin (1% v/v, Thermo Fisher Scientific) and 2-mercaptoethanol (0.1 % v/v, Thermo Fisher Scientific). CT26_β2m^-/-^ and RMA-S were cultured in RPMI 1640 medium (Thermo Fisher Scientific) supplemented with Fetal Bovine Serum (FBS) (10 % v/v, Thermo Fisher Scientific), HEPES (1 % v/v, Thermo Fisher Scientific), sodium pyruvate (1 % v/v, Thermo Fisher Scientific), penicillin/streptomycin (1 % v/v, Thermo Fisher Scientific) and 2-mercaptoethanol (0.1 % v/v, Thermo Fisher Scientific). U87 and HepG2 cells were cultured in EMEM medium (Thermo Fisher Scientific) supplemented with Fetal Bovine Serum (FBS) (10 % v/v, Thermo Fisher Scientific), HEPES (1 % v/v, Thermo Fisher Scientific), sodium pyruvate (1 % v/v, Thermo Fisher Scientific), penicillin/streptomycin (1 % v/v, Thermo Fisher Scientific) and 2-mercaptoethanol (0.1 % v/v, Thermo Fisher Scientific).

#### Mice

Six- to eight-week-old female CD45.2^+^Thy1.2^+^ C57BL/6 (C57BL/6J), BALB/c (BALB/cByJ) mice and CD45.1^+^ mice (B6.SJL-*Ptprc^a^ Pepc^b^*/BoyCrl) were purchased from Charles River Laboratories (Lyon, France). CD45.2^+^CRISPR-Cas9 Knock-in mice (Gt(ROSA)26Sor^tm1.1(CAG-cas9*,-EGFP)Fezh/J^) were purchased from the Jackson Laboratory and maintained at the EPFL’s pathogen-free facility. All mice were housed in the EPFL Center of PhenoGenomics and were kept in individually ventilated cages, at 19-23 °C, with 45-65 % humidity, and with a 12 h dark/light cycle. Experimental procedures in mouse studies were approved by the Swiss authorities (Canton of Vaud, animal protocol ID 3206, 3533, 3902, 3912, 3915, and 3009) and performed under the guidelines from the EPFL Center of PhenoGenomics. Female NOD-Prkdc^scid^ Il2rg^em1^/Smoc (M-NSG) mice were purchased from Shanghai Model Organisms Center, Inc. and housed at the specific pathogen-free conditions according to the guidelines for experimental animals at the University of Science and Technology of China. Experimental procedures were approved by the Ethics Committee at the University of Science and Technology of China (USTCACUC192201040).

#### Human blood samples

PBMC were collected from healthy donors at the Blood Center of Anhui Province (Hefei, China). Human PBMCs were received from healthy volunteers with written informed consent and the protocol was approved by the Ethics Committee of the University of Science and Technology of China (Hefei, China, 2019-N(H)-121).

### METHOD DETAILS

#### IL-21−Fc protein production and purification

The IL-21−Fc fusion protein containing a mouse or human IL-21, fused at the N-terminal with a noncytolytic IgG Fc. It was expressed by HEK293-E cells (Thermo Fisher Scientific) at the EPFL Protein Expression Core Facility. The supernatant of culture medium containing IL-21−Fc fusion protein was harvested by centrifugation after 7 days and was filtered through a 0.22-μm membrane to remove cell debris and obtain a clear solution. A HiTrap Protein A affinity chromatography column (Cytiva, 17-0403-01, 5 ml) on AKTA pure 25 (GE Healthcare) was used to capture the recombinant IL-21−Fc from the clarified, filtered expression supernatant. IL-21−Fc was eluted with elution buffer (0.05 M sodium citrate, 0.3 M NaCl, pH 3.0). The eluted protein was immediately collected into the neutralization buffer (1.0 M Tris–HCl, pH 10.0) followed by concentrating by 10 kDa membrane ultrafiltrating (GE Healthcare). The concentrated protein solution was further purified with Superdex 200 increase size-exclusion chromatography (GE Healthcare) at a flow rate of 1.0 mL/min with PBS on AKTA pure 25 (Figure S2A). The purified protein was aliquoted and stored at −80 °C for subsequent usage. The purity of IL-21−Fc was confirmed with sodium dodecyl sulfate polyacrylamide gel electrophoresis (SDS-PAGE, Figure S2B) and the bioactivity was evaluated by comparing to commercial IL-21 (Biolegend) (Figure S2C).

#### Preparation of NK cells

Spleens from Thy1.2^+^CD45.2^+^ C57BL/6 or BALB/c mice were mechanically disrupted and gridded through a 70 μm strainer (Fisher Scientific, Pittsburgh, PA, USA). NK cells were isolated by magnetic-activated cell sorting (MACS) from mouse splenocytes with the NK Cell Isolation Kit (Miltenyi Biotec). Isolated mouse NK cells were cultured for 5 days in the complete RPMI 1640, which contained FBS (10 % v/v), HEPES (1 % v/v), penicillin/streptomycin (1 % v/v), sodium pyruvate (1 % v/v), 2-mercaptoethanol (0.1 % v/v) and supplemented with mouse IL-2 (50 ng/mL, Biolegend) to maintain NK cells survival and growth. PBMC were collected from healthy donors at the Blood Center of Anhui Province (Hefei, China). Human NK cells were isolated by CD56 Microbeads, human (Miltenyi Biotec), followed by activating and culturing in complete RPMI 1640 medium containing hIL-2 (100 U/mL) and human IL-15 (20 ng/mL) for 24 h.

#### Preparation of WT and LDHA KO NK cells

NK cells were isolated from splenocytes of CRISPR–Cas9 knock-in mice. To generate WT NK and LDHA KO NK cells, NK cells were isolated in the same method described above and activated with IL-2 (50 ng/mL) contained complete RPMI-1640 medium for 24 hours, followed by transducing with retrovirus containing scramble control guide RNA or LDHA-targeting gRNA on 6-well plate coated with protamine (10 µg/mL, Thermo Fisher Scientific). Transduced NK cells were then cultured and expanded for an additional 2 days prior to use. The pools of gRNAs targeting LDHA (LDHA1: TTGGTGAGTATACCTTGGGC, LDHA2: GGGACTTGGTGAGTATACCT, LDHA3: GGGCAGGTTACAATGACACA, LDHA4: GCACCGGCTAAAGTCACTGT) and a scramble control gRNA (TGCGAATACGCCCACGCGAT) were designed using the public available gRNA design tool^76^.

#### *In vitro* co-culture of NK cells and tumor cells

Tumor cells were cultured and maintained in the corresponding complete medium as described. Harvested tumor cells were resuspended and seeded in the 24-well plate in complete medium at 37°C for 12-16 h, then aspirate the tumor culture medium and adding activated NK cells resuspension prepared as described, IL-21−Fc were supplemented to the complete RPMI 1640 medium at 100 ng/mL. After 24 hours of co-culture, the NK cells phenotype and function were determined by flow cytometry. To determine the killing efficacy towards target cells, tumor cells were seeded in the 96-well flat-bottom microplate and cultured as described above. NK cells were added at indicated effector-to-target ratio in the absence or presence of IL-21−Fc (100 ng/mL). Tumor cell viability was measured by LDH cytotoxicity assay (Thermo Fisher Scientific) after 5 hours of co-culture.

#### LDHA KO NK cell transfer

Thy1.2^+^CD45.2^+^ mice were inoculated with B16F10_ β2m^-/-^ tumor cells (8 × 10^5^, s.c.) and received sublethal irradiation on day 6. Mice received intravenously adoptive transfer of LDHA KO NK cells (1 × 10^6^) or WT NK cells (1 × 10^6^) on day 7 followed by injection of IL-21−Fc (20 µg, i.t.) or PBS every other day until day 13. On day 14, mice were sacrificed and tumor-infiltrating leukocytes (TILs) were analyzed by flow cytometry.

#### Antitumor therapy and rechallenging experiments

Mice bearing established tumors with palpable area around 20-40 mm^2^ (day 7 post inoculation or as indicated) were treated with injection of IL-21−Fc or PBS (i.t.) every other day, 8 doses in total as indicated. For the combination of IL-15SA and IL-21−Fc, mice bearing established tumors with the palpable area around 20-40 mm^2^ (day 7 post inoculation or as indicated) were treated with injection of IL-15SA (5 µg, i.t.) on day 7 and day 14, followed by injection of IL-21−Fc (20 µg, i.t.) or PBS every other day from day 7, 8 doses in total as indicated. Tumor area and body weight were measured every other day. For the treatment of the xenografted HepG2 tumor model, mice were treated with adoptive transfer of activated hNK cells intravenously (5 × 10^6^), followed by injection of hIL-2 (75 kU, i.p.), and injection of hIL-21−Fc (20 µg, i.t.) or PBS every other day from day 7, 6 doses in total. Tumor area was calculated according to Area = Length × Width from caliper measurements of 2 orthogonal diameters. Mice were euthanized upon body weight loss beyond 15 % of pre-dosing weight, or the tumor area reached 150 mm^2^, or the animal had become moribund (according to the animal license). For tumor rechallenge studies, CT26_β2m^-/-^ (1 × 10^5^, s.c.), and B16F10_β2m^-/-^ (1 × 10^5^, s.c.) were implanted into the left flanks of cured mice from treatment groups on day 90 post primary tumor inoculation. Age-matched wild-type mice were inoculated (s.c.) with the same number of tumor cells as control. Survival of rechallenge mice was monitored for other 60 days.

#### Selectively depletion of specific immune cell subsets

C57BL/6J mice were inoculated with B16F10_β2m^-/-^ tumor cells (8 × 10^5^, s.c.) and started treatment from day 7. Depletion antibodies anti-mouse CD8α (Clone 2.43, BioXcell), anti-mouse CD4 (Clone YTS191, BioXcell), or anti-mouse NK1.1 (Clone PK136, BioXcell) were injected (400 ug, i.p.) by day 5 that two days prior to treatment for every three days. Peripheral blood was collected by day 12 that post three times injections of depletion antibodies, blood cells resuspension was prepared followed by staining with indicated antibodies and analyze by flow cytometry (Attune NxT Flow Cytometer, Invitrogen / Thermal Fischer Scientific).

#### Analysis of tumor-infiltrating immune cells

BALB/c mice were inoculated with CT26_β2m^-/-^ tumor cells (8 × 10^5^, s.c.) and received treatment on day 7 post tumor inoculation. Mice were received four doses of administration of IL-21−Fc (20 µg, i.t.) or PBS control every two days starting from day 7. On day 14, tumors were dissected and weighed. The dissected tumors were mechanically minced and digested through stirring in RPMI 1640 medium that contained collagenases Type IV (1 mg/mL, Thermo Fisher Scientific), dispase II (100 μg/mL, Sigma-Aldrich), hyaluronidase (100 μg/mL, Sigma-Aldrich), and DNase I (100 μg/mL, Sigma-Aldrich) at 37 °C for 60 min. Red blood cells were removed with ACK lysing buffer after digestion. TILs were enriched by density gradient centrifugation against Percoll (GE Healthcare) and resuspended in FACS buffer (1 × PBS containing 0.2 % BSA (wt / v), Sigma-Aldrich), then stained with indicated antibodies and analyzed with flow cytometry (Attune NxT Flow Cytometer).

#### Flow cytometry analysis

Cells were blocked with anti-mouse CD16/32 antibody at 4°C for 20 min at first, followed by incubation with indicated antibodies 4 °C for another 20 min. After surface staining, cells were stained by live/dead dye 4’,6-diamidino-2-phenylindole (DAPI, Sigma-Aldrich) or Zombie Aqua Fixable Dye (BioLegend). Cells then were washed and resuspended with FACS buffer for flow cytometry analysis. For intracellular cytokines staining, cells were previously stimulated by a Cell Stimulation Cocktail (protein transport inhibitors included, Thermo Fisher Scientific) in advance at 37°C for 4-6 h. Next, cells were stained for surface markers and or Zombie Aqua Fixable Dye as described above. Cells then were fixed and permeabilized with a Cytofix/Cytoperm™ Fixation/Permeabilization Solution Kit (BD Biosciences) according to the manufacturer’s instructions. For transcriptional factors staining, cells were first stained for surface markers and or Zombie Aqua Fixable Dye as described above. Next, cells were fixed and permeabilized with a Foxp3/Transcription Factor Staining Buffer Set (eBioscience) for transcription factor staining according to the manufacturer’s instructions. Data was collected using an Attune NxT Flow Cytometer.

#### Cell sorting

Tumor-infiltrating CD3^-^NKp46^+^ cells from CT26_β2m^-/-^ tumors, and splenic CD3^-^NKp46^+^ cells from tumor-bearing mice were firstly enriched by density gradient centrifugation against Percoll (GE Healthcare), followed by MACS using mouse NK isolation kit (Miltenyi Biotec), and then stained with surface markers and DAPI followed by sorting with an Aria II sorter (BD Biosciences) or SONY SH800S (Sony Biotech) at the EPFL Flow Cytometry Core Facility.

#### Seahorse assay

Seahorse assay was performed to measure the OCR and ECAR of NK cells. *In vitro* cultured NK cells or sorted NK cells from tumors with different treatments were seeded in a seahorse cell culture plate (Seahorse Bioscience) at the density of 3 × 10^5^/well in the non-CO_2_ incubator at 37 °C for 40 min. During the seahorse assay, cells were treated with oligomycin (1 μM, Sigma-Aldrich), Carbonyl cyanide-4-(trifluoromethoxy) phenylhydrazone (FCCP, 2 μM, Sigma-Aldrich), rotenone (0.5 μM, Sigma-Aldrich), antimycin A (0.5 μM, Sigma-Aldrich), glucose (10 mM, Sigma-Aldrich), and 2-Deoxy-D-glucose (2-DG, 50 mM, Sigma-Aldrich). Each condition was performed with 3–6 replicates in a single experiment. OCR and ECAR were measured by an XF96 Seahorse Extracellular Flux Analyzer (Seahorse Bioscience) following the manufacturer’s instructions. OXPHOS and glycolysis were calculated according to previous reports^77^.

#### Metabolic inhibitor treatments

NK cells were prepared as previously outlined. Subsequently, they were cultured in complete RPMI 1640 medium supplemented with IL-2 (50 ng/mL), in the presence or absence of IL-21−Fc (100 ng/mL), alongside specified metabolic inhibitors (2-DG at 0.5 mM, Oligomycin at 1 µM, FX II at 10 µM) for a duration of 24 hours. Flow cytometry was employed to analyze the cells.

#### Bulk RNA-seq and bioinformatics analysis

Mice were inoculated with CT26_β2m^-/-^ tumor cells (8 × 10^5^, s.c.). On day 14, mice were sacrificed and TILs were prepared as described above. CD45.2^+^CD3^-^NKp46^+^ NK cells were sorted as described above. Total RNA extraction was carried out using the RNeasy Mini Kit (Qiagen), followed by quality assessment using TapeStation 4200 (Agilent Technologies) to ensure RNA integrity (scores > 7). High-quality RNA samples were then stored at −80°C until library preparation. Libraries for mRNA-seq were prepared utilizing the NEBNext Ultra II Directional RNA Library Prep with PolyA method (NEB), starting from 20 ng RNA, in accordance with the NEBNext Ultra II Directional RNA Library Prep Kit, Version 4.0_4/21 (internal exp code CSX063). Each library, distinguished by unique dual indexes, was subsequently loaded onto a NovaSeq 6000 flow cell (Illumina) and sequenced following the manufacturer’s recommendations, yielding at least 40 million pairs of 60 nucleotides-long reads per sample. Raw reads underwent adapter trimming using BCL convert v3.9.3 (Illumina) and quality assessment using fastQC v0.11.9. Alignment and quantification of all RNA sequencing data were conducted using the STAR and Salmon methods via the nf-core v3.9 pipeline. Alignment was performed against the mouse genome GRCm38. Read counts were normalized for library size using the TMM method from EdgeR and Limma-Voom. Differential expression analysis was carried out using limma, after filtering out genes with an average Transcripts Per Million (TPM) less than two. Pathway analysis was performed by GSEA with cluster profiler and mSigDB using Hallmark, KEGG, Reactome, GO Biological Process, and Wiki Pathways.

#### scRNA-seq and bioinformatics analysis

Mice were inoculated with CT26_β2m^-/-^ tumor cells (8 × 10^5^, s.c.), followed by administration of IL-15SA (5 µg, i.t.) on day 7, and 4 doses administration of IL-21−Fc (20 µg, i.t.) or PBS control every other day. On day 14, mice were sacrificed and TILs were prepared as described above. CD45.2^+^CD3^-^NKp46^+^ NK cells were sorted as described above, cells from the same treatment group were pooled together and resuspended at 3 × 10^5^ cells/mL in PBS. Next, 100 µL cell resuspension was loaded onto a standard GEXSCOPE® microwell chip. scRNA-seq library was constructed using the GEXSCOPE® Single-Cell RNA Library Kit according to the manufacturer’s instructions (Singleron Biotechnologies GmbH). Individual libraries were diluted to equimolar and pooled for sequencing. Pools were sequenced on a lane of an S4 flow cell on an Illumina NovaSeq 6000 with 150 bp paired-end reads to a total depth of 90 GB per sample and an average read depth per cell of 50,000 reads. Raw reads were processed with FastQC to evaluate the quality of raw data.

Reads were demultiplexed according to cell barcodes and unique molecular identifiers (UMI), and poly-A tails and adaptor sequences were removed by Cutadapt using CeleScope v1.13.0. After quality control, reads were mapped to the reference genome (Mus_musculus_ensembl_92). Gene counts and UMI counts were acquired via feature Counts software. Gene expression matrix files for subsequent analyses were generated based on gene counts and UMI counts. Downstream data analysis was performed with the Seurat v4 pipeline and Scanpy pipeline^78^. Cells were first filtered based on three metrics: 1) the number of detected genes per cell between 500 to 5000; 2) the proportion of mitochondrial gene counts (UMIs from mitochondrial genes / total UMIs) must be less than 10 %; 3) the total counts of cells must between 1000 to 10000. Subsequently, the gene expression data was log-normalized utilizing the Seurat “NormalizeData” function from Seurat. No significant batch effects were observed between the two samples. Principal component analysis (PCA) was conducted on the scaled data using the top 800 highly variable genes (HVGs) identified by “FindVariableFeatures”. Notably, genes related to programs and compartments were excluded based on gene sets from the SignatuR package (https://github.com/carmonalab/SignatuR). The gene expression matrix was further reduced to 15 principal components (PCs), as well as to 2 UMAP dimensions for visualization. Unsupervised clusters were calculated on the PC space using the “FindClusters” function from Seurat at a resolution of 0.2. Differentially expressed genes (DEGs) were identified using the “rank_genes_groups” function from Scanpy for pairwise comparison between the two conditions on the log-transformed counts data. The top 1500 significant genes were selected to perform GSEA using the Python package “gseapy”^79^, with the annotation gene set obtained from the “enrichr” database^80^. To re-cluster cells based on the list of KEGG metabolism pathway, 7 categories encompassing 53 related pathways were selected from the KEGG database. The “score_genes” function was employed to calculate the metabolism score for each cell in each pathway, using these cores to define unsupervised clusters. The ratio of observed to expected, representing the enrichment of cell clusters in the two conditions^81^, was calculated based on the chi-square test, where the 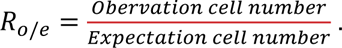

#### Metabolite extraction and protein quantification

Mice were inoculated with CT26_β2m^-/-^ tumor cells (8 × 10^5^, s.c.), followed by administration of IL-15SA (5 µg, i.t.) on day 7, and 4 doses administration of IL-21−Fc (20 µg, i.t.) or PBS control every other day. On day 14, mice were sacrificed and TILs were prepared as described above. CD45.2^+^CD3^-^NKp46^+^ NK cells were sorted as described above, cells from the same treatment group were pooled together and resuspended at 5 × 10^5^ cells/mL in 50 µL PBS. Resuspended cells were pre-extracted and homogenized by the addition of 200 µL of MeOH, in the Cryolys Precellys 24 sample Homogenizer (2 × 20 seconds at 10000 rpm, Bertin Technologies, Rockville, MD, US) with ceramic beads. The bead beater was air-cooled down at a flow rate of 110 L/min at 6 bars. Homogenized extracts were centrifuged for 15 minutes at 4000 g at 4°C (Hermle). The resulting supernatant was collected and evaporated to dryness in a vacuum concentrator (LabConco). Dried sample extracts were resuspended in MeOH: H_2_O (4:1, v/v) according to the total protein content. The protein pellets were evaporated and lysed in 20 mM Tris-HCl (pH 7.5), 4M guanidine hydrochloride, 150 mM NaCl, 1 mM Na_2_EDTA, 1 mM EGTA, 1 % Triton, 2.5 mM sodium pyrophosphate, 1 mM beta-glycerophosphate, 1 mM Na_3_VO_4_, 1 µg/ml leupeptin using the Cryolys Precellys 24 sample Homogenizer (2 × 20 seconds at 10000 rpm, Bertin Technologies) with ceramic beads. BCA Protein Assay Kit (Thermo Scientific) was used to measure (A562nm) total protein concentration (Hidex, Turku, Finland).

#### Multiple pathway targeted analysis

Extracted samples were analyzed by Hydrophilic Interaction Liquid Chromatography coupled to tandem mass spectrometry (HILIC - MS/MS) in both positive and negative ionization modes using a 6495 triple quadrupole system (QqQ) interfaced with 1290 UHPLC system (Agilent Technologies). In positive mode, the chromatographic separation was carried out in an Acquity BEH Amide, 1.7 μm, 100 mm × 2.1 mm I.D. column (Waters). The mobile phase was composed of A = 20 mM ammonium formate and 0.1 % formic acid (FA) in water and B = 0.1 % FA in acetonitrile (ACN). The linear gradient elution from 95 % B (0-1.5 min) down to 45 % B was applied (1.5 min −17 min) and these conditions were held for 2 min. Then initial chromatographic condition was maintained as a post-run for 5 min for column re-equilibration. The flow rate was 400 μL/min, column temperature 25 °C, and sample injection volume 2 µl. ESI source conditions were set as follows: dry gas temperature 290 °C, nebulizer 35 psi and flow 14 L/min, sheath gas temperature 350 °C and flow 12 L/min, nozzle voltage 0 V, and capillary voltage 2000 V. Dynamic Multiple Reaction Monitoring (DMRM) was used as acquisition mode with a total cycle time of 600 ms. Optimized collision energies for each metabolite were applied. In negative mode, a SeQuant ZIC-pHILIC (100 mm, 2.1 mm I.D. and 5 μm particle size, Merck, Damstadt, Germany) column was used. The mobile phase was composed of A = 20 mM ammonium Acetate and 20 mM NH_4_OH in water at pH 9.7 and B = 100 % ACN. The linear gradient elution from 90 % (0-1.5 min) to 50 % B (8-11 min) down to 45 % B (12-15 min). Finally, the initial chromatographic conditions were established as a post-run during 9 min for column re-equilibration. The flow rate was 300 μL/min, column temperature 30 °C, and sample injection volume 2 µl. ESI source conditions were set as follows: dry gas temperature 290 °C and flow 14 L/min, sheath gas temperature 350 °C, nebulizer 45 psi, and flow 12 L/min, nozzle voltage 0 V, and capillary voltage −2000 V. Dynamic Multiple Reaction Monitoring (dMRM) was used as acquisition mode with a total cycle time of 600 ms. Optimized collision energies for each metabolite were applied. Raw LC-MS/MS data was processed using the Agilent Quantitative analysis software (version B.07.00, MassHunter Agilent technologies). Relative quantification of metabolites was based on EIC (Extracted Ion Chromatogram) areas for the monitored MRM transitions. The obtained tables (containing peak areas of detected metabolites across all samples, Table S2) were exported to “R” software http://cran.r-project.org/ and signal intensity drift correction and noise filtering (if necessary, using CV (QC features) > 30 %) was done within the MRM PROBS software. The preprocessed data with peak areas were imported into Metaboanalyst 5.0 for further data analysis.

#### Ethics statement

Experiments and handling of mice were conducted under federal, state, and local guidelines with approval from the Swiss authorities (Canton of Vaud, animal protocol ID 3206, 3533, 3902, 3912, 3915, and 3009) and performed in accordance under the guidelines from the EPFL Center of PhenoGenomics. Human PBMCs were received from healthy volunteers with written informed consent and the protocol was approved by the Ethics Committee of the University of Science and Technology of China (Hefei, China, 2019-N(H)-121). All NSG mouse experiments were approved by the Institutional Animal Care and Use Committee of the University of Science and Technology of China (Hefei, China, USTCACUC192201040).

### QUANTIFICATION AND STATISTICAL ANALYSIS

#### Statistical analysis

Statistical analysis was performed using GraphPad Prism 10 (GraphPad software). Data are presented as mean ± s.e.m. unless indicated. Comparisons of the two groups were performed by using a two-tailed paired or unpaired Student’s t-test. Comparisons of multiple groups at a single time point were performed by using one-way analysis of variance (ANOVA) and Tukey’s test. Survival data were analyzed using the Log-rank test. No statistically significant (NS) differences were considered when P-values were larger than 0.05.

