## Supplementary information for "Metabolic reinvigoration of NK cells by IL-21 enhances immunotherapy against MHC-I deficient solid tumors"

### IL-21 reprograms NK cell metabolism to enhance antitumor immunity against solid tumors

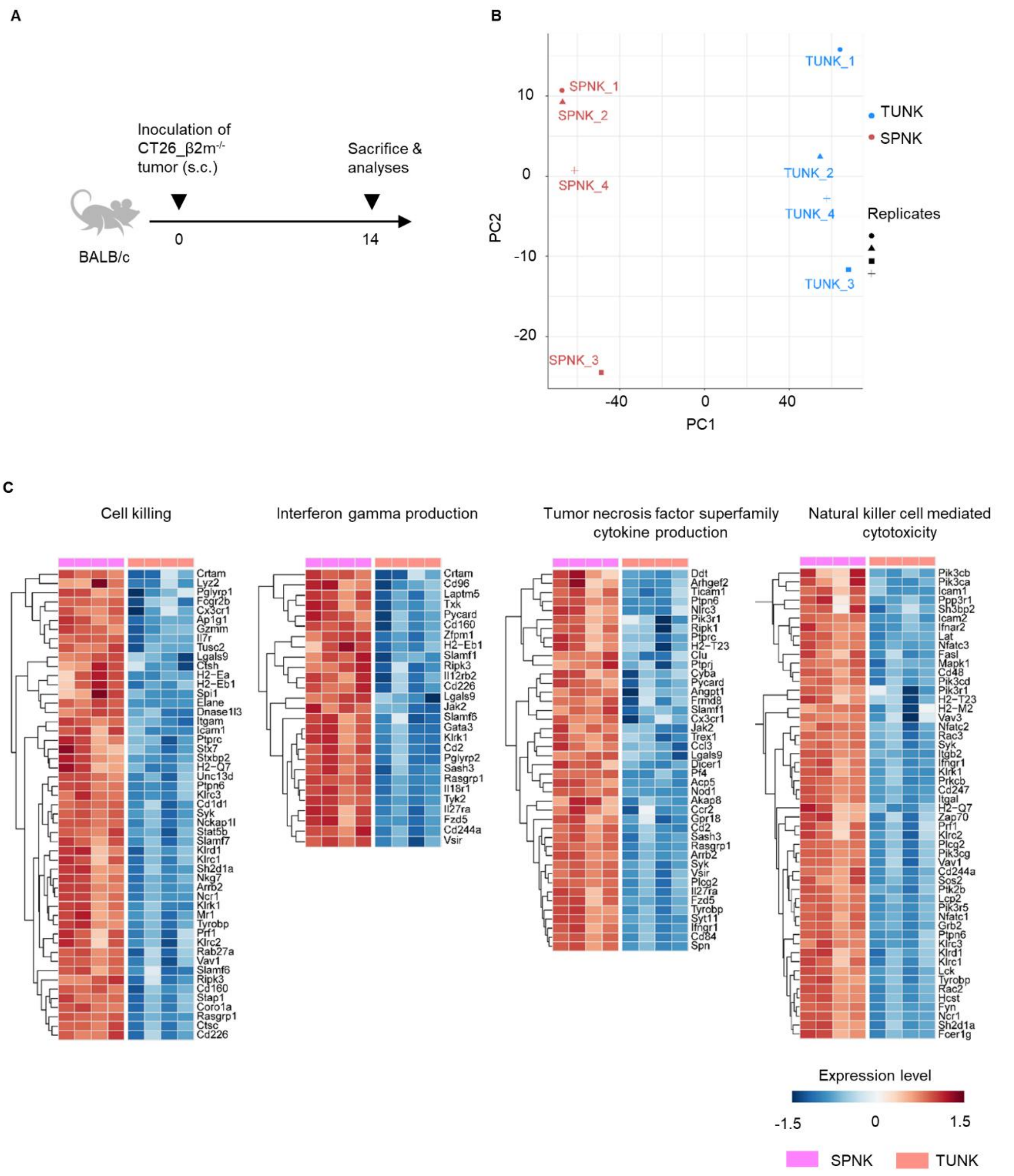


**Figure S1. Transcriptomic analysis of NK cells in spleen and tumor, related to Figure 1.**

(A) Experimental timeline for Figure 1A-D and Figure 1E-I.

(B) The principal component analysis (PCA) of 50 % most highly varying genes of SPNK and TUNK. Counts were normalized for library size using TMM method from EdgeR and Limma-Voom. Differential expression of 9836 genes was computed with limma after filtering out genes with an average TPM (Transcripts Per Million) less than two.

(C) Heatmap of differentially expressed genes ranked by GSEA between SPNK and TUNK in Figure 1F-I.

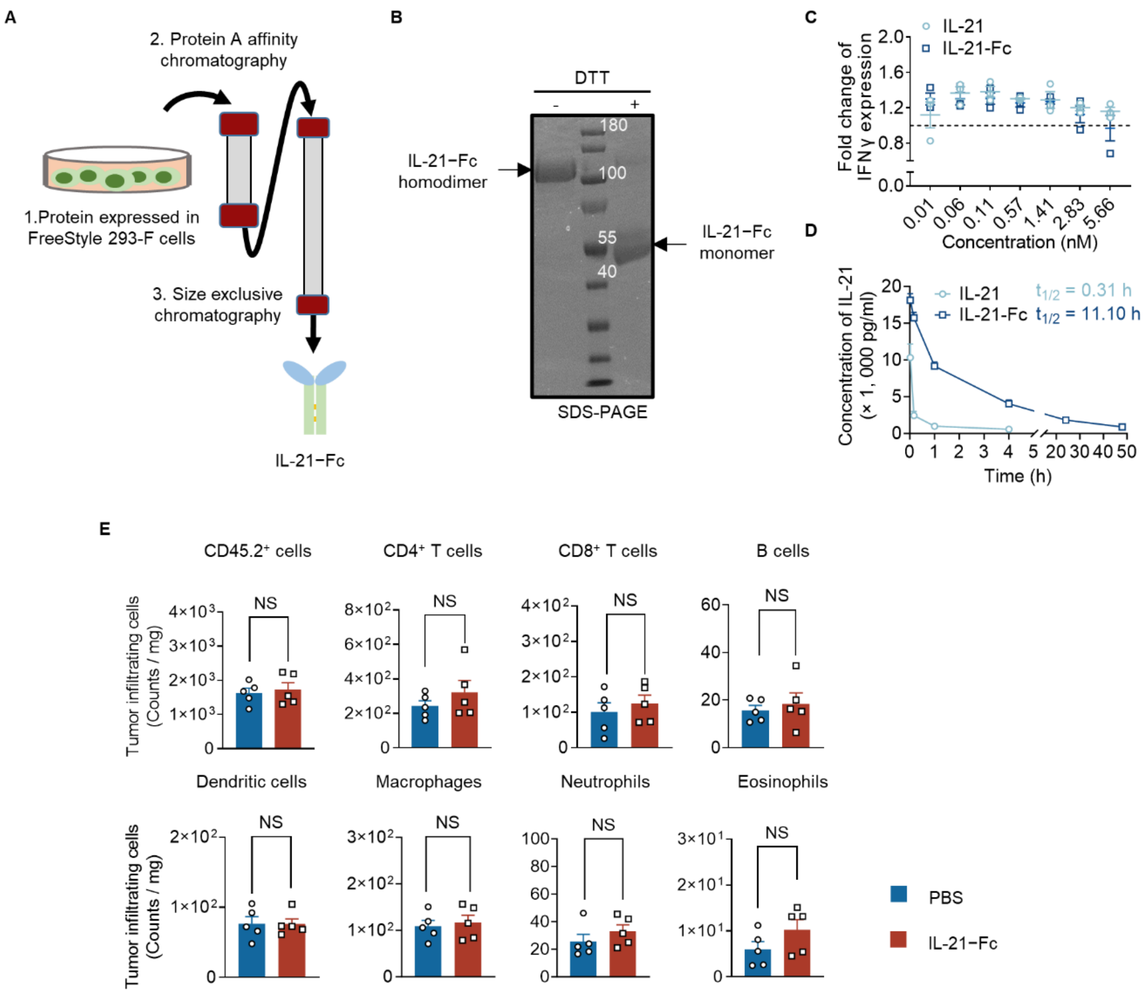


**Figure S2. IL-21−Fc has negligible effect on other tumor-infiltrating immune cells in tumor, related Figure 2.**

(A) Scheme of the production and purification of IL-21−Fc fusion protein. IL-21−Fc was expressed by HEK293-E cells. IL-21−Fc containing supernatant was firstly captured with HiTrap Protein A affinity chromatography and further purified with Superdex 200 increase size-exclusion chromatography.

(B) SDS-PAGE analysis of purified IL-21−Fc. DTT, dithiothreitol.

(C) NK cells activation is similar as described in Figure A-B. Activated NK cells were cultured in the presence of IL-21 or fusion IL-21−Fc at a gradient concentration for 24 h. Shown is the fold changes of IFNγ expression that normalized by the PBS group.

(D) The pharmacokinetics in plasma and half-life of IL-21 or IL-21−Fc (n = 5 animals).

(E) The counts of various tumor-infiltrating immune cells in CT26_β2m$^{-/-}$ tumors (n = 5 animals). Also see Figure 2E. All data represent the mean ± s.e.m. and are analyzed by two-sided Student's t-test. NS, not significant (P > 0.05).

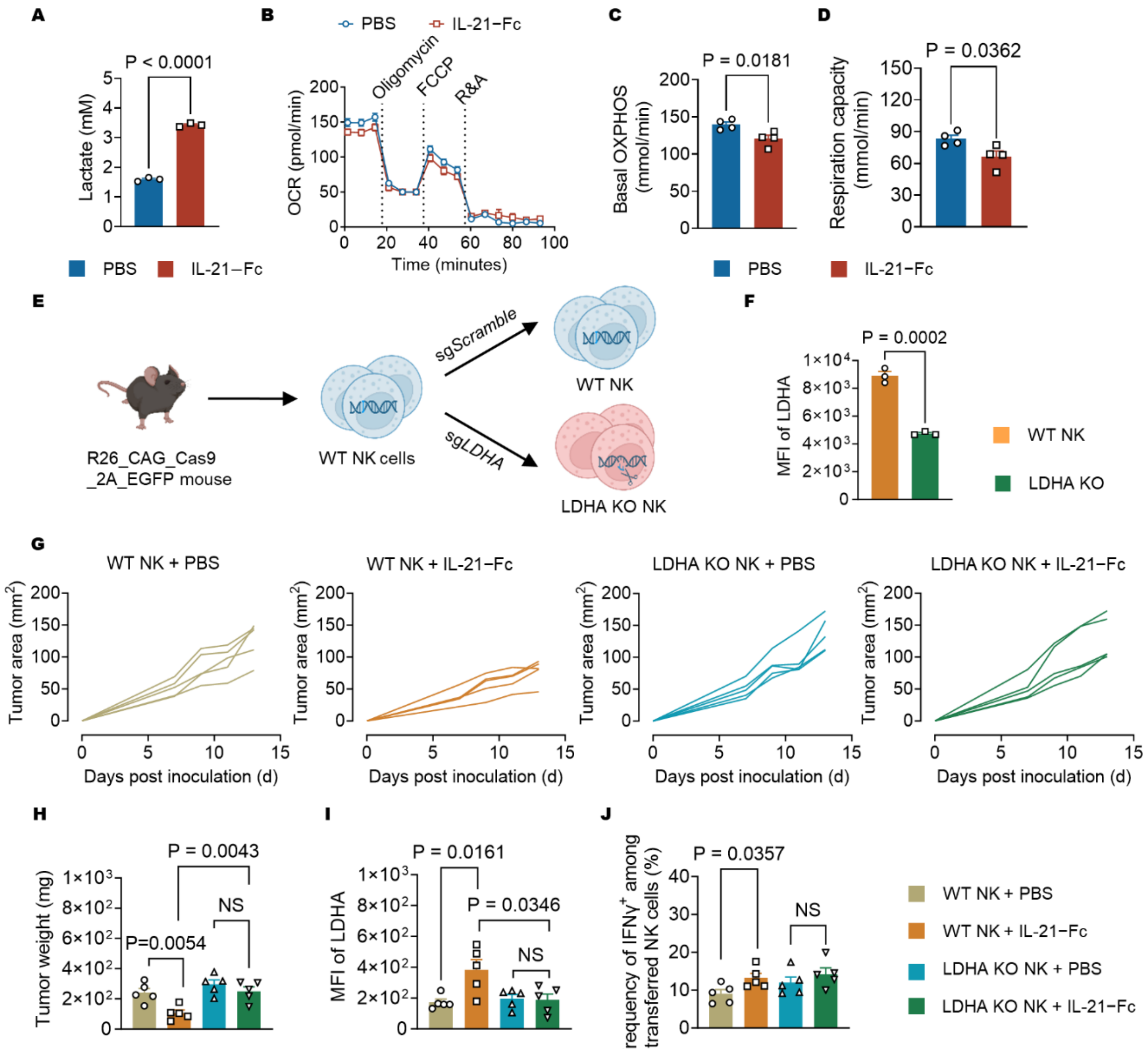


**Figure S3. IL-21−Fc reprograms NK cell metabolism by elevating LDHA-dependent glycolysis, related to Figure 3.**

(A) The extracellular lactate concentrations in the supernatant of NK cells.

(B) The real-time OCR curves of NK cells (n = 4 replicates).

(C) The basal OCR of NK cells.

(D) The maximum OCR of NK cells.

(E) Schematic illustration of LDHA KO NK cells generation.

(F) The MFI of LDHA of WT NK cells and LDHA KO NK cells.

(G-J) The experiment setting was described in Figure 3O.

(G) The individual tumor growth curves.

(H) Tumor weight on day 14 post inoculation.

(I) MFI of LDHA of transferred NK cells.

(J) The frequency of IFNγ$^{+}$ NK cells among transferred NK cells.

All data represent the mean ± s.e.m. and are analyzed by two-sided Student's t-test (**A-D, F**) or one-way ANOVA with Tukey's test (**H-J**).

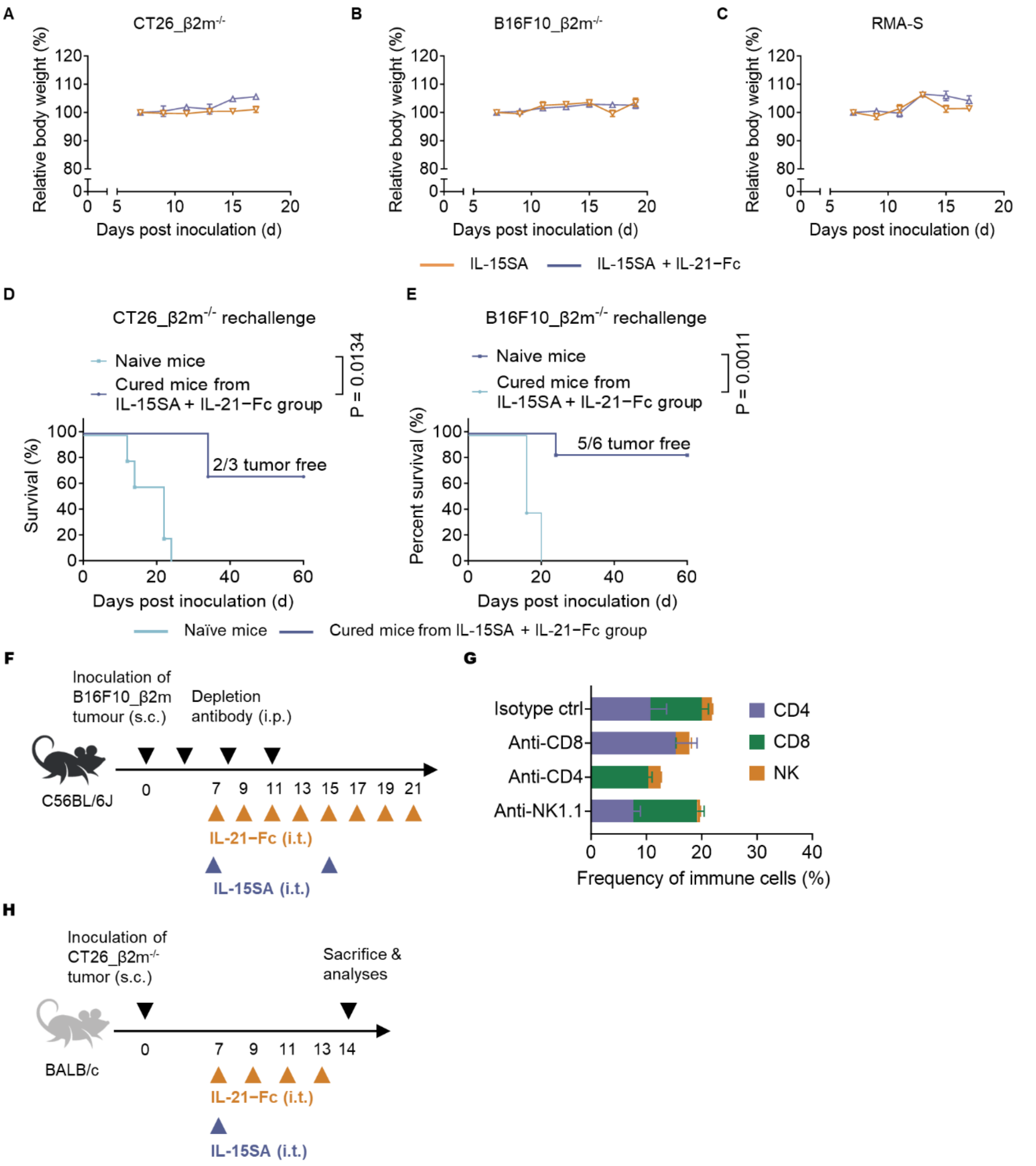


**Figure S4. NK cells are indispensable for the enhanced antitumor efficacy mediated by combinatory therapy of IL-21−Fc and IL-15SA, related to Figure 4.**

(A-E) The experimental setting was described in Figure 4A-G.

(A-C) The body weight changes of mice with CT26_ β2m$^{-/-}$ tumors (A), B16F10_β2m$^{-/-}$ tumors (B), and RMA-S tumors (C)

(D-E) Kaplan-Meier survival curves of naïve or cured mice rechallenged with CT26_β2m$^{-/-}$ cells (D), or B16F10_β2m$^{-/-}$ cells (E).

(F) Experiment timeline that descried in Figure 4H.

(G) The frequency of various immune cells in blood one day post administration (i.p.) of immune cell depletion antibodies.

(H) The experiment timeline described in Figure 4I-J.

All data represent the mean ± s.e.m. and are analyzed by Log-rank test for survival curves (D-E).

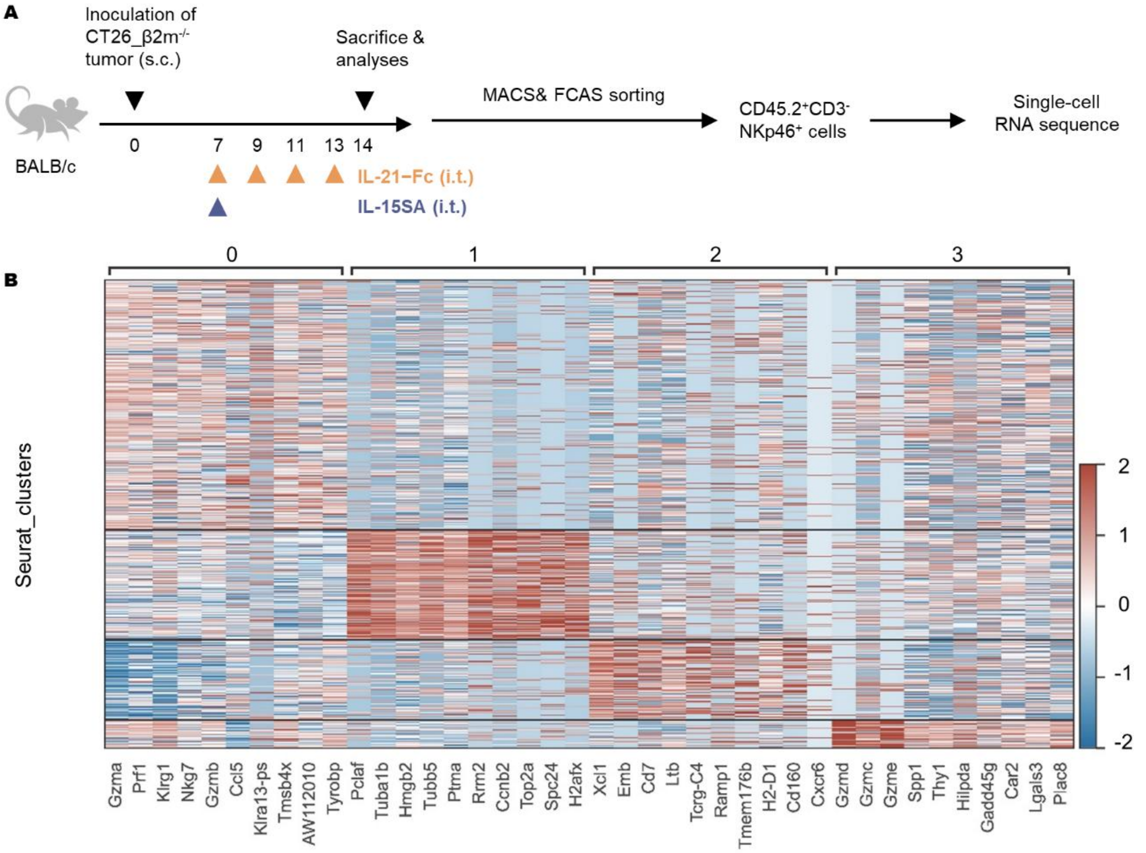


**Figure S5. Single-cell transcriptional profiling of tumor-infiltrating NK cells treated with IL-21−Fc, related to Figure 5.**

(A) The experiment timeline for Figure 5A-B.

(B) Heatmap of the top expressed gene within the four clusters defined in Figure 5B.

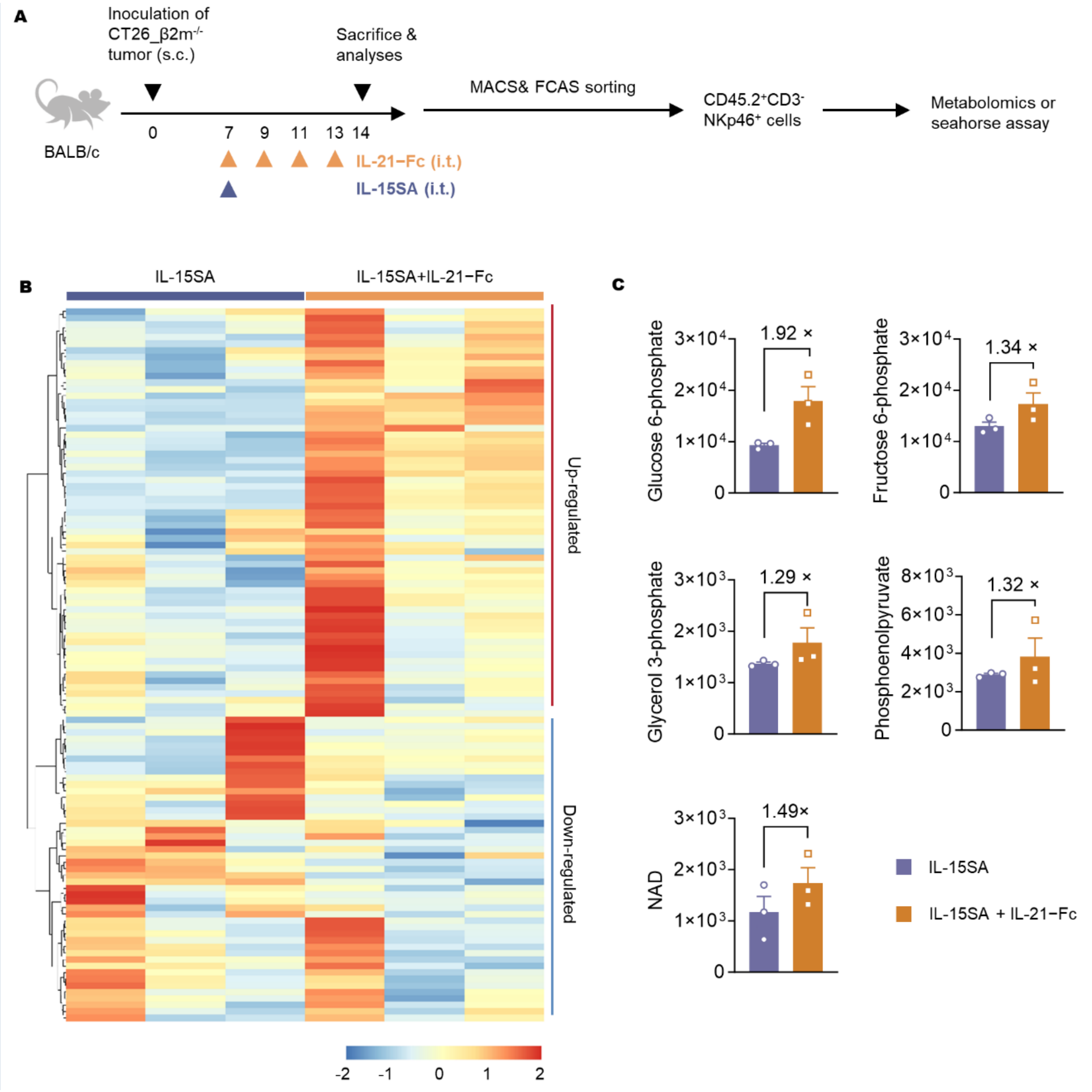


**Figure S6. IL-21−Fc programs the metabolic activity of tumor-infiltrating NK cells for elevated glycolysis, related to Figure 6.**

(A) The experiment timeline of Figure 6E-I.

(B) Differential metabolites expression of NK cells treated with IL-15SA compared to IL-15SA + IL-21−Fc. Shown are heatmap of the differential abundance of detected metabolites.

(C) Relative quantification of representative metabolites involved in the glycolysis pathway. All data represent the mean ± s.e.m. and are analyzed by a two-sided Student's t-test.

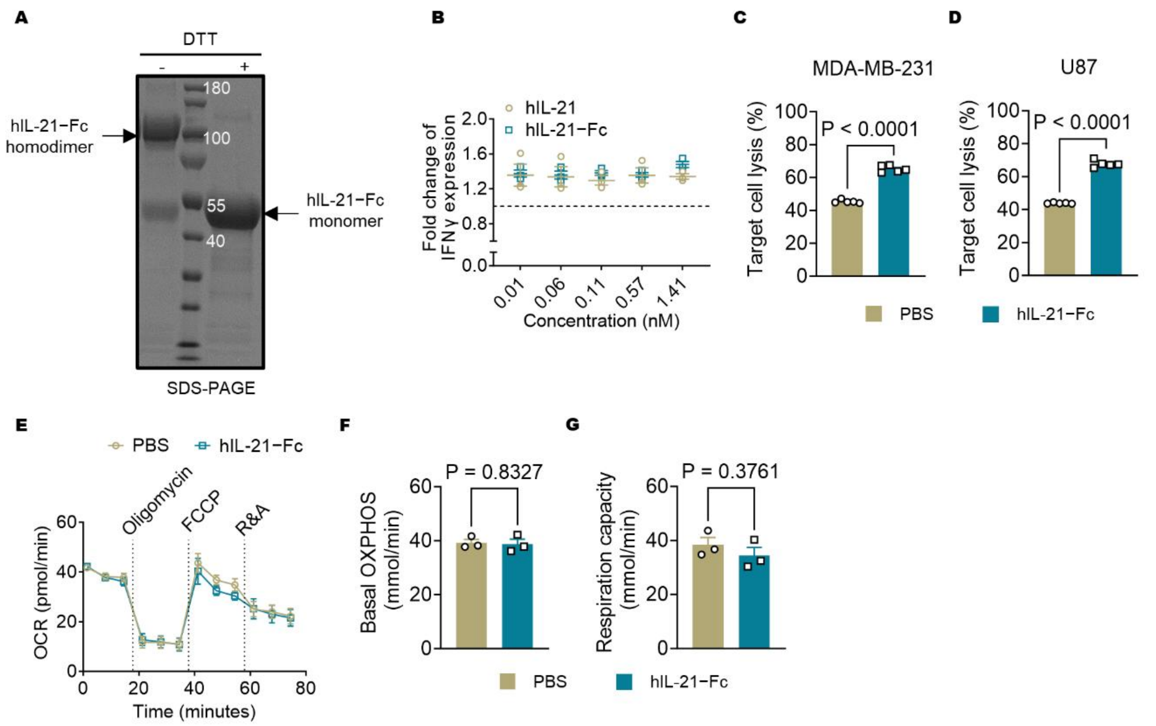


**Figure S7. Human IL-21−Fc enhances cytotoxicity of NK cells, related to Figure 7.**
(A) The production of hIL-21−Fc was similar as described in Figure S2A. Shown is the SDS-PAGE of purified hIL-21−Fc.
(B) hNK cells were activated as described in Figure 7A-D and cultured in the presence of hIL-21 or fusion hIL-21−Fc at a gradient concentration. Shown is the fold change of IFNγ expression of hNK cells normalized by that in the PBS group.
(C-D) hNK cells were co-cultured with cancer cells in the presence of or absence of hIL-21−Fc (100 ng/mL) at an effector-to-target (E/T) ratio of 0.5 for 5 h.
(C) The percentage of lysis of MDA-MB-231 breast cancer cells.
(D) The percentage of lysis of U87 glioblastoma cells.
(E) The real-time OCR curves (n =3 replicates).
(F) The basal OCR of NK cells.
(G) The maximum OCR of NK cells.
All data represent the mean ± s.e.m. and are analyzed by a two-sided Student’s t-test.

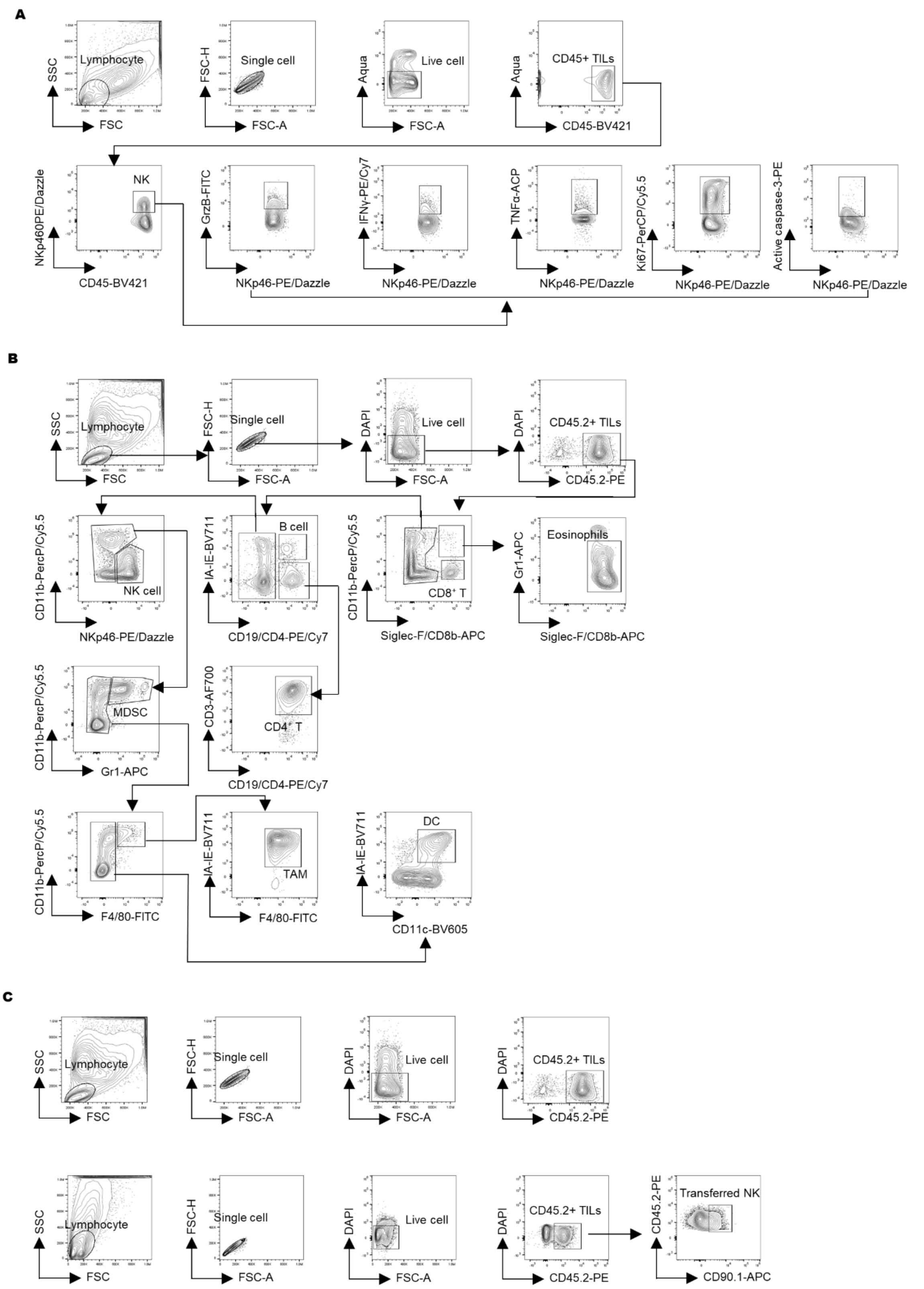


**Figure S8. Flow cytometry gating strategies.**

(A) Representative flow cytometry gating strategies, related to Figure 1A, Figure 2F-J, Figure 4I and J.

(B) Representative flow cytometry gating strategies related to Figure S2E.

(C) Representative flow cytometry gating strategies related to Figure 3Q and Figure S3H-J.
